# Blood prognostic biomarker signatures for hemorrhagic cerebral cavernous malformations (CCMs)

**DOI:** 10.1101/2023.07.01.547354

**Authors:** Jacob Croft, Brian Grajeda, Johnathan S Abou-Fadel, Cameron C Ellis, Estevao Igor, Igor C. Almeida, Jun Zhang

## Abstract

**Background:** Cerebral cavernous malformations (CCMs) is a neurological disorder that causes enlarged intracranial capillaries in the brain, leading to an increased risk of hemorrhagic strokes, which is a leading cause of death and disability worldwide.

**Objectives:** The current treatment options for CCMs are limited, highlighting the need for prognostic biomarkers to predict the risk of hemorrhagic events to better inform treatment decisions and identify future pharmacological targets.

**Eligibility Criteria:** The study is centered on a comparative proteomic analysis between hemorrhagic CCMs (HCs) and healthy controls, while excluding patients with non-hemorrhagic CCMs (NHCs) from the analysis due to the experimental design.

**Sources of Evidence:** Recent research has identified several serum biomarkers and blood circulating biomarkers in a selected cohort of homogeneous CCM patients and animal models.

**Method:** Proteomic profiles from both human and mouse CCM models were examined, and pathway enrichment analyses were performed using three approaches (GO, KEGG, and DOSE). To account for multiple comparisons, t-tests were employed to evaluate differences. A p-value below 0.05 was deemed statistically significant.

**Results:** The authors have developed the first panel of candidate biomarker signatures, featuring both etiological and prognostic markers in two distinct pathways. This panel of biomarker signatures demonstrates a robust correlation with the likelihood of hemorrhagic CCMs.

**Conclusions:** This groundbreaking biomarker panel paves the way for further investigation of potential blood biomarkers to determine the risk of hemorrhagic CCMs.

## Introduction

Cerebral cavernous malformations (CCMs) is a neurological disorder that causes enlarged intracranial capillaries in the brain, leading to an increased risk of hemorrhagic strokes ^1, 2^. This condition is particularly prevalent in the Hispanic population, with the highest rates observed in individuals with the CCM1 gene mutation ^3–6^. The common Hispanic BACA-CCM1 mutation has been traced back to several generations of descendants in the West Texas borderland area, which also has the highest percentage of Hispanics in the United States and is considered the epicenter for CCMs in North America by stroke specialists. Unfortunately, the majority of individuals with the CCM gene mutation are largely asymptomatic, but when symptoms do occur, the disease has often reached the stage of focal hemorrhage, leading to significant morbidity ^7–11^.

Currently, the treatment options for CCMs are limited to neurosurgical removal for most lesions or gamma knife radiosurgery for deep stem CCM lesions ^12–15^. These treatments are invasive and can have serious consequences for patients, highlighting the need for prognostic biomarkers to predict the risk of hemorrhagic events to better inform treatment decisions and identify future pharmacological targets. Recent research has identified several etiological serum biomarkers and blood circulating biomarkers in a selected cohort of homogeneous CCM patients sharing the BACA-CCM1 mutation ^7^. Based on the previously identified biomarkers, the authors have developed the first panel of candidate biomarkers to predict the risk of hemorrhagic CCMs in the largest recorded sample size from local Hispanic CCM patients. This biomarker panel has the potential to significantly improve patient outcomes and reduce the morbidity associated with this debilitating condition.

Biomarkers are an essential tool in predicting, screening, diagnosing, prognosticating, or stratifying risk for disease outcomes. They can be cellular, histological, molecular, physiological, or radiographic characteristics and can be used alone or as a panel with multiple targets (FDA-NIH BEST resources). Blood biomarkers, especially as prognostic biomarkers, are advantageous due to their low cost, feasibility, and acceptability for diagnostic and prognostic applications. They have been long sought as diagnostic and prognostic tools for various disorders, mainly ischemic strokes ^16–21^. Thus far, the progress in developing biomarker-driven prediction models for hemorrhagic CCMs has been restricted. A handful of initial investigations have been carried out in human and animal models, yet they have faced considerable constraints^7, 22–25^.

In this research, we have discovered the inaugural set of biomarker signatures closely linked to hemorrhagic risk. This innovative biomarker panel lays the groundwork for further assessment of potential blood biomarkers in determining the risk of hemorrhagic CCMs. This could pave the way for a reformed approach incorporating updated clinical definitions and substantially enhance the management of hemorrhagic strokes.

## Materials and Methods

The objective of this study was to identify a set of blood-based biomarkers that can predict the prognosis of different stages of hemorrhagic strokes. To accomplish this, we began an exploratory project with the hypothesis that prognostic blood biomarkers for hemorrhagic risk, discovered within a genetically well-defined familial CCM cohort, can be expanded and applied to sporadic CCM cases and ultimately to a broader range of hemorrhagic strokes. Moreover, ongoing debates and unresolved issues persist concerning the underlying causes of both familial and sporadic CCM forms, especially whether they originate from familial CCM mutations or other unknown genes ^2, 26–30^. The future validation of potential biomarkers stemming from this research will undoubtedly help to address this query.

Since this is a comprehensive investigation of CCM1 mutation effects, and serves as a basis for further large-scale studies, the participants were chosen based on their homogenous genetic predisposition and symptomatic presentation to establish the differential protein expression patterns of CCM1 mutations compared to that of age-/gender-matched control groups. We are confident to analyze the experimental outcomes with a cost-effective model for protein expression data to minimize type 1 errors (type-1 errors=0.05) in the cost of sacrificing for type-2 error (missing some potential targeted proteins) as detailed in statistical analysis section ^31^.

In this study, we carried out a proteomics analysis involving a cohort of human patients with familial cerebral cavernous malformations (fCCM) resulting from a uniform BACA CCM1 hemizygous mutation, alongside their age- and gender-matched healthy controls (n=14). In addition, we examined Ccm1 mutant mice along with their wild-type (WT) counterparts (n=6) to strengthen our analytical capabilities. While this Omics research substantially deviates from traditional epidemiological analysis, it is important to note that this investigation is conducted within the context of a clinical trial focused on biomarker identification and validation. As a result, we have rigorously adhered to the STROBE (Strengthening the Reporting of Observational Studies in Epidemiology) guidelines^32^ throughout the process of preparing this manuscript.

### 1. fCCM patient cohorts’ recruitment procedure

The criteria for the International Classification of Diseases (ICD) codes 9/10 must be met in order for a patient’s medical history to be considered for the study on CCMs (categorized as 228.00, 228.02, 228.09/Q28.3, D18.00, D18.02). Neurovascular disorders (747.81/Q28.2, Q04.9, G93.9) and codes related to hemorrhagic stroke and epilepsy (430, 431, 432.1, 432.9, and 345.00, 345.01/I60.9, I61.9, I62.00, I62.9 and G40.A01, G40.A09, G40.A11, G40A.19) may also be used as supplementary criteria. The authorized IRB protocol permits the enrollment and consent of participants ranging from 8 to 89 years old. Considering the substantial Hispanic population in the vicinity, the study primarily focuses on including minority individuals, although it does not impose any restrictions, as detailed in. Individuals with CCMs can be classified into two distinct groups: non-hemorrhagic CCMs (NHCs) and hemorrhagic CCMs (HCs). For this comparative proteomic study, only hemorrhagic CCMs (HCs) and healthy controls (Ctrls) were utilized.

### 2. Data collection

Data for the study was collected through structured interviews conducted in person by clinical co-investigators and stored on secure PCs. The correlation between blood biomarker levels and disease severity in fCCM patients were analyzed using statistical methods. Odds ratios and 95% confidence intervals were calculated using stratified data analysis and logistic regression to determine the relationship between the blood levels of biomarkers and the risk of hemorrhagic stroke in fCCM patients. Comparisons were made between symptomatic fCCM carriers (HCs) and healthy controls to determine the correlation between blood levels of biomarkers and the odds of hemorrhagic stroke. All statistical tests were two-tailed with a significance level of p<0.05.

### 3. Biomarker data collection from proteomics

To discover new biomarkers through high-throughput omics methods using human serum samples from fCCM patients and healthy matched controls, we conducted a comprehensive and impartial search. Our optimized procedures include: *<u>3-1) Abundant serum protein depletion</u>*: We processed 10 serum samples from hemorrhagic fCCM patients with BACA-CCM1 mutations and matched controls for proteomic analysis ^33–36^.. The total protein concentration was determined using the Bicinchoninic Acid (BCA) assay (Bio-Rad). Then, using High Select Mini Spin Columns (Thermo Scientific), we removed highly abundant proteins such as albumin, immunoglobulins, fibrinogen, and transferrin, to detect low-abundant biomarkers. The protein concentration was re-measured using the BCA assay. *<u>3-2) Protein Enrichment:</u>* The serum samples were processed with a ProteoMiner kit (Bio-Rad) and the protein concentration was determined using the BCA and fluorescence assays. *<u>3-3) Trypsin Digestion and iTRAQ Labeling</u>*: The serum proteins were reduced, alkylated, desalted, and buffer-changed, and then digested with trypsin and labeled with iTRAQ reagents. The labeled peptides were subjected to a Phoenix Peptide Cleanup Kit to remove excess labeling and salt, and were lyophilized and stored at −80°C. *<u>3-4) Raw Proteomic Data Acquisition:</u>* All proteomic data were acquired through tandem mass spectrometer (MS/MS) analysis, including NanoLC-MS/MS analysis, using a Q Exactive Plus Hybrid Quadrupole-Orbitrap Mass Spectrometer (Thermo Scientific).

By following these optimized procedures, we aimed to identify new biomarkers in fCCM patients compared to healthy matched controls through high-throughput omics utilizing human serum samples.

### 4. Biomarker identification through pathway analyses with bioinformatics tools

The Bioconductor R package was employed carry out biological process analysis and functional interaction network analysis ^37–39^. The groupGO function in clusterProfiler of the Bioconductor was used to determine the functional profile of the pathway components by grouping genes based on their gene ontology ^40–43^. A vector of UniProt Accession numbers was provided for the gene argument, and the Bioconductor Genome-Wide Annotation for Mouse was used as the database. The keytype argument was set to “UNIPROT,” converting the UniProt Accession numbers into Entrez Gene IDs, and the level was set to GO levels 2. This process was repeated for all three sub-ontologies and followed by hierarchical clustering analysis. Complementing this process the enrichGO function in clusterProfiler of the Bioconductor further explored the grouping of the gene ontology creating pathways visualizing functional profiles of where these proteins were located in the previously mentioned sub-ontology levels. Additionally, GO, KEGG and DOSE pathways, modules, and pathway enrichment analysis were performed using Bioconductor and its associated functions^40, 44–50^. The enrichDO function in clusterProfiler of the Bioconductor was as a systems approach to analyze the disease ontology from the Entrez Gene ID mapping related disease based off of shared genes ^50^. Finally all pathway enrichments were completed analyzing the Entrez Gene Id to determine the molecular functions and relationship shared throughout the subject.

### 5. Statistical analysis

Significance was adjusted for multiple comparisons, and differences between groups were compared using the Student’s t-test or paired samples t-test. A p-value less than 0.05 was considered statistically significant. Our analysis indicated that our proposed sample size would be suitable for exploring protein expression differences between CCM1 deficient and normal groups (HCs and age-/gender-matched Ctrls). Results were visualized using ggplot2 with log2fc for fold changes ^51^.

A power analysis was employed to assess the ability of the sample sizes to yield significant findings. By setting a significance level of 0.05 to reduce type 1 errors, the power level dropped below 80%, making the study more prone to type 2 errors, or false negatives. To counteract this, only participants with a homogenous genetic background were recruited, ensuring a population with a known inheritance history. The selection of subjects was further refined by considering phenotypical manifestations in a clinical setting, which guaranteed a sample with actively expressed proteins relevant to the condition. Consequently, instead of relying on a power analysis-based population, a cost-effective model was adopted, resulting in statistically significant findings that contribute to the field and the potential identification of biomarkers.

## Results

### 1. Differentially expressed serum proteins were identified through proteomics in two species

Using proteomics approach, we identified a total of 69 confirmed serum proteins from 189 peptide entries in humans, and 105 confirmed serum proteins from 168 peptide entries in mice. By comparing the distribution of expressed protein profiles between CCM1-deficient subjects and normal controls across species, we found that there were more down-regulated DEPs than up-regulated DEPs in both humans (Fig. 1-A) and mice (Fig. 1-B). Of these, 35 genes were found to be uniquely expressed and shared between the two species (Fig. 1-C). Comparative heatmaps visualize the differential expression of serum proteins (DEPs) between CCM1-deficient subjects and normal controls in both species (Fig. 1-D). This suggests that the DEPs shared by both species could serve as potential blood circulating biomarkers for hemorrhagic CCMs. We observed a congruous trend of DEPs in both species, which was further highlighted in the Volcano plots in both humans (Fig. 1-E) and mice (Fig. 1-F). The data presented showcases the involvement of DEPs in several biological processes, making it an excellent foundation for the following gene pathway and enrichment analysis.

**Figure 1.**
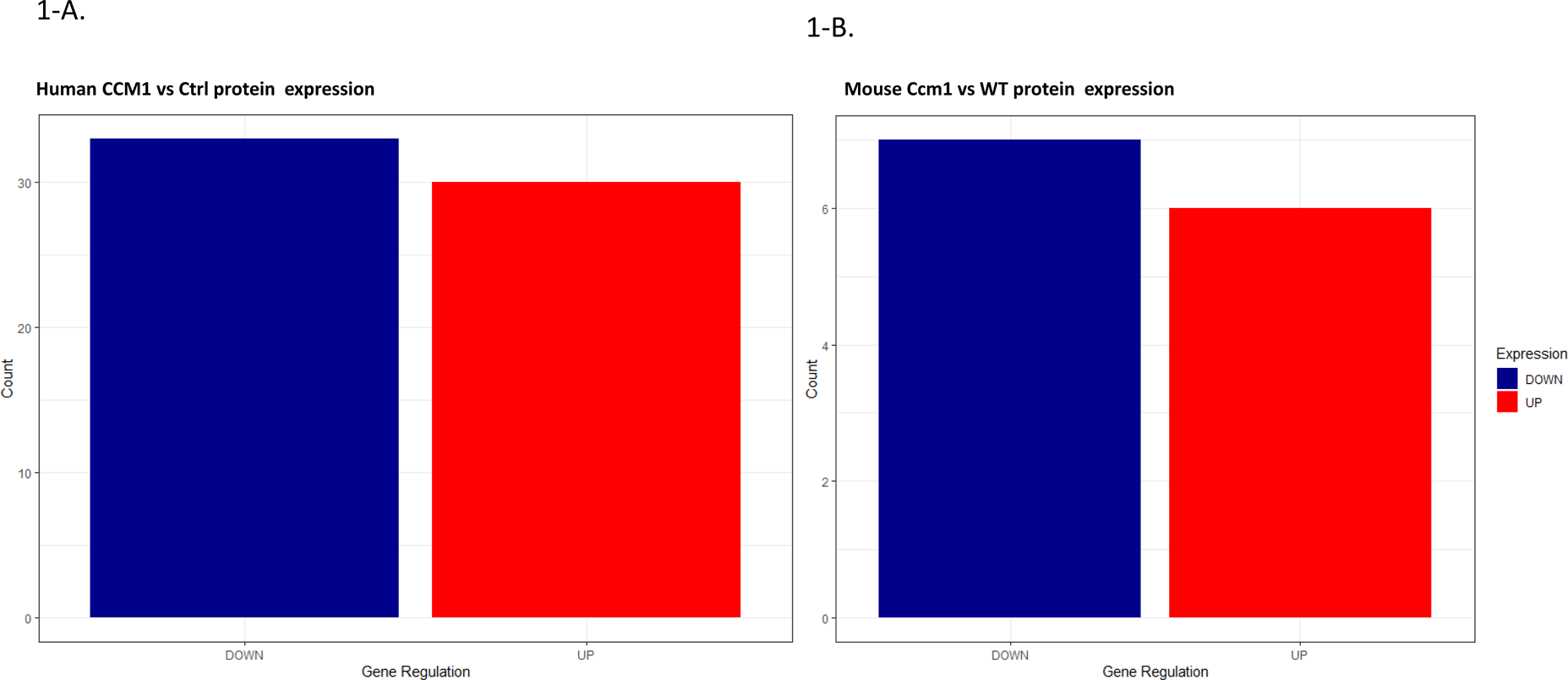
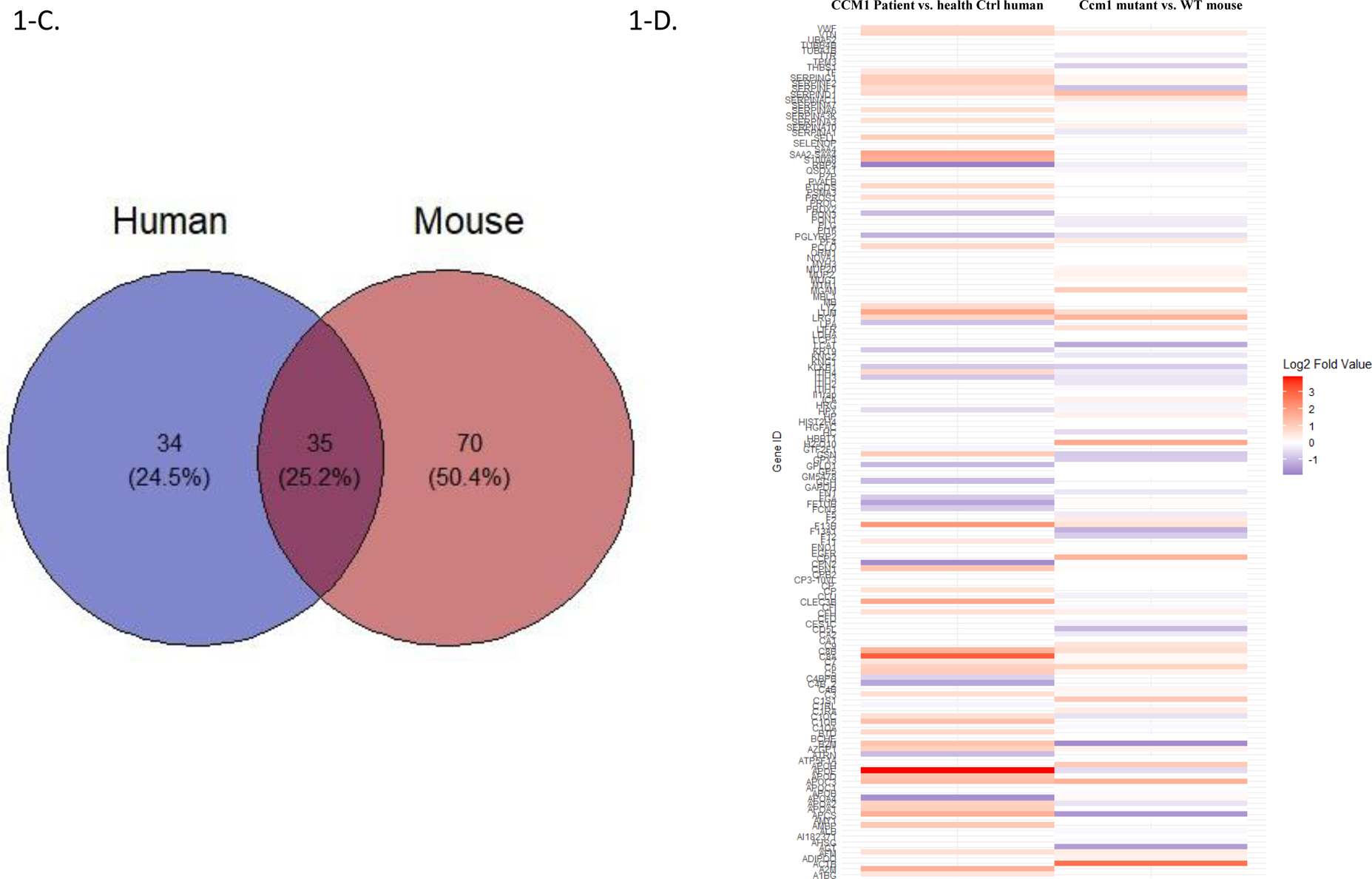
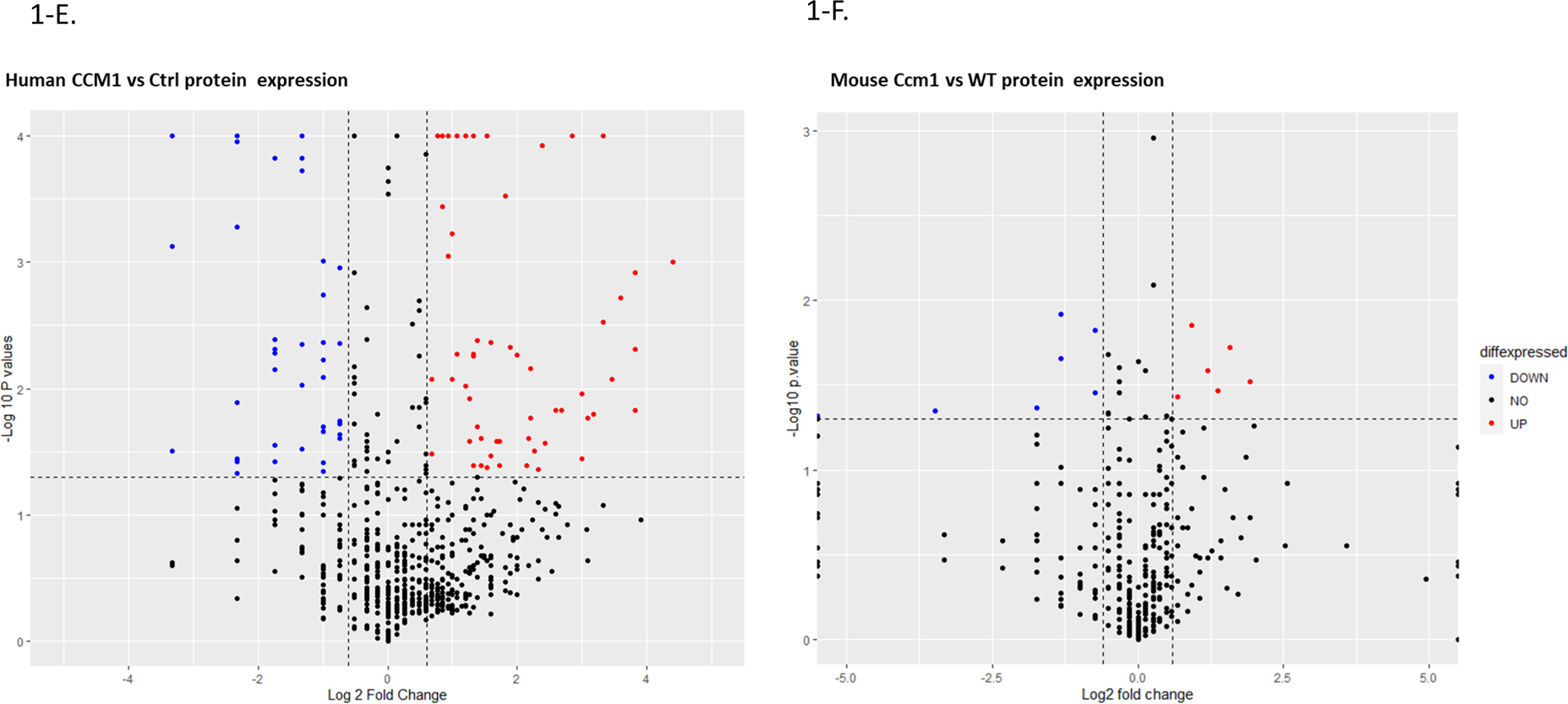
Differentially expressed serum proteins (DEPs) identified in CCM1-deficient subjects and normal controls across species by utilizing high-throughput comparative proteomics. Statistical significance of differentially expressed serum proteins (DEPs) were identified by analyzing the distribution of expressed protein profiles. A-B. Distribution of highly expressed serum proteins in both CCM1-deficient subjects and normal controls across species. The distribution of DEPs was overlaid by comparing CCM1-deficient subjects to normal controls in both human (A) and mice (B). Up-(red bar) and down-(blue bar) regulation were determined by fold change values. C-D. Unique patterns of the DEPs were identified by comparing CCM1-deficient individuals and normal controls in two species. A Venn diagram displays unique and co-expressed proteins between humans and mice (C). Comparative heatmaps illustrate protein expression changes between CCM1-deficient subjects and controls in both species (D). The heatmaps displayed protein expression values for each related protein on the Y-axis. E-F. Volcano plots display DEPs between CCM1-deficient subjects and healthy controls. This visualization highlights proteins present in both control and confirmed CCM-deficient subjects, demonstrating how the CCM1 protein expression varies in comparison to controls. A p-value, derived from a student’s t-test, is shown in a −log10 function relative to the log2 fold change. This approach helps regulate infinite fold values for unique values, preventing distortion of the graph. The data points within the plot represent fold changes in protein expressions for CCM1-deficient subjects compared to the control group in both humans (E) and mice (F). To identify more significant results, the values of up- and down-regulated proteins from figures E and F were further refined using a −log10 p-value threshold.

### 2. CCM1-associated signaling pathways were identified through pathway analyses among DEPs in two species

Gene pathway and enrichment analysis is a widely used method in omics to identify overrepresented gene sets, such as pathways, gene ontology terms, or disease-associated genes, in a specific subset of genes or proteins. This approach provides valuable insights into the underlying biological processes of a disease phenotype. In our study, we conducted gene ontology (GO), disease ontology (DO), and KEGG pathway enrichment analyses to comparatively examine the functional characteristics and biological pathways of the differentially expressed proteins (DEPs) identified in our study.

To determine the statistical significance of the biological functions within the group of DEPs identified in our study, we firstly conducted group gene ontology (GO) and pathway analysis with systematic functional annotation^52–54^. Given the significant number of DEPs identified in both species, we used these DEPs for the initial biological profiling through functional interaction network and pathway analyses. GO enrichment analysis was performed based on the described method ^55^. Enriched GO pathway dotplots offer a comparative visualization of functional enrichment outcomes in both human (Fig. 2-A, Suppl. Figs. 1A, 1C, 2A, 2C, 2E, 2F) and mouse (Fig. 2-B, Suppl. Figs. 1B, 1D, 2B, 2D, 2G, 2H) datasets, enabling us to discern patterns in extensive biological data and assess the differentially expressed proteins (DEPs) affected by CCM1 deficiency.

**Figure 2.**
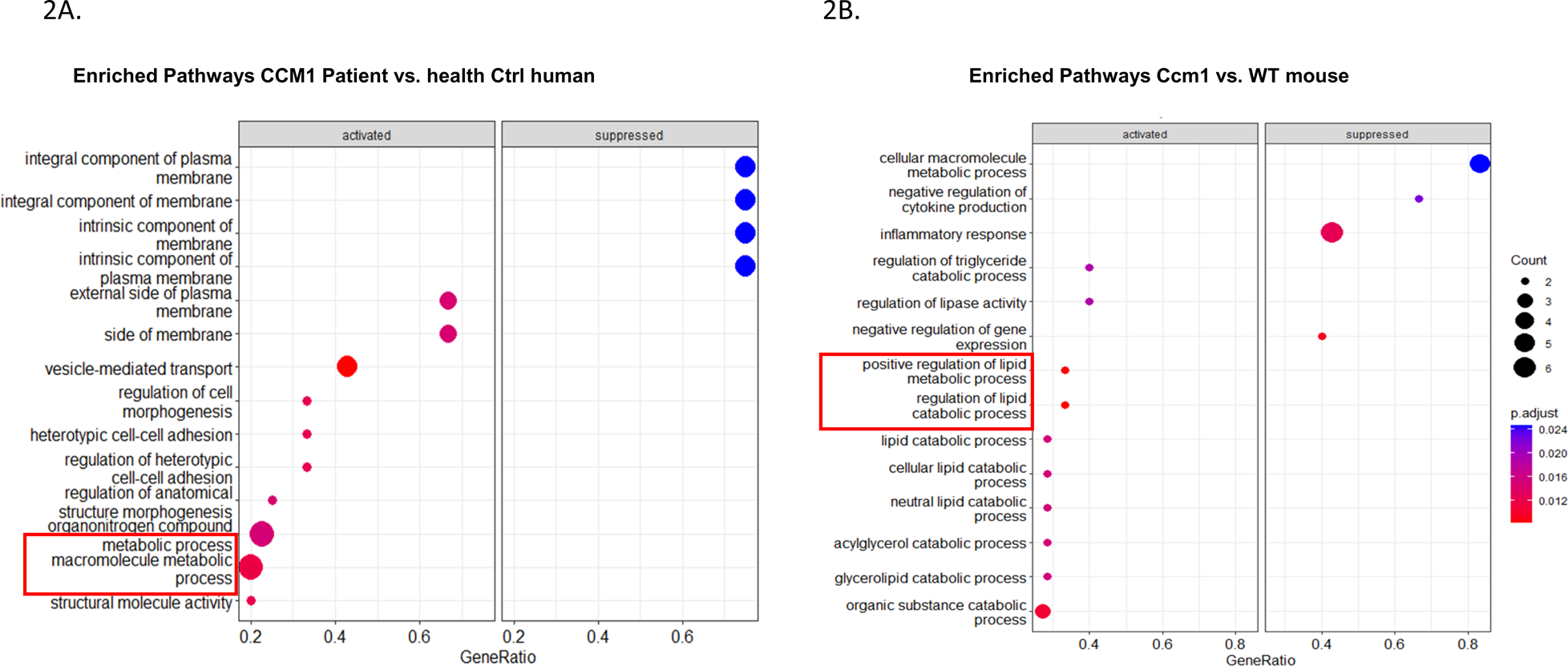
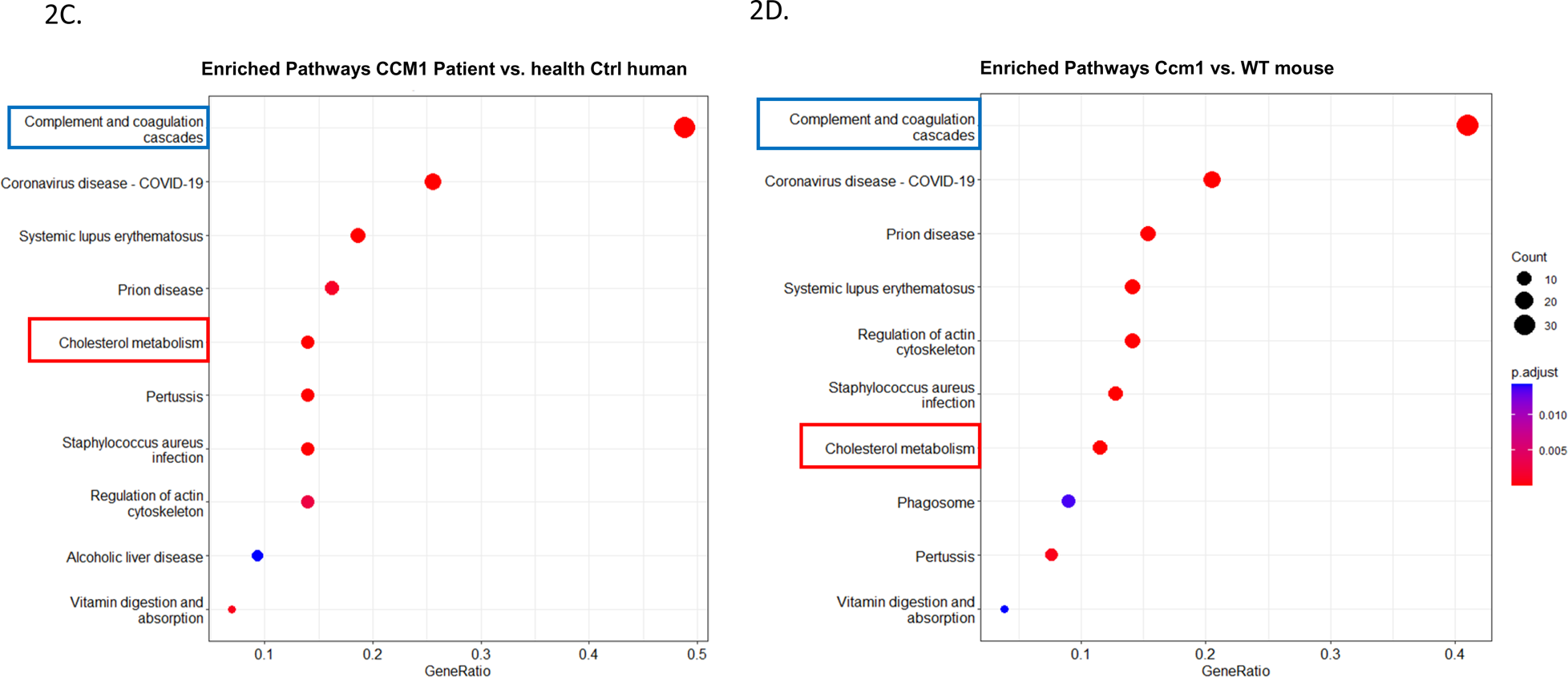
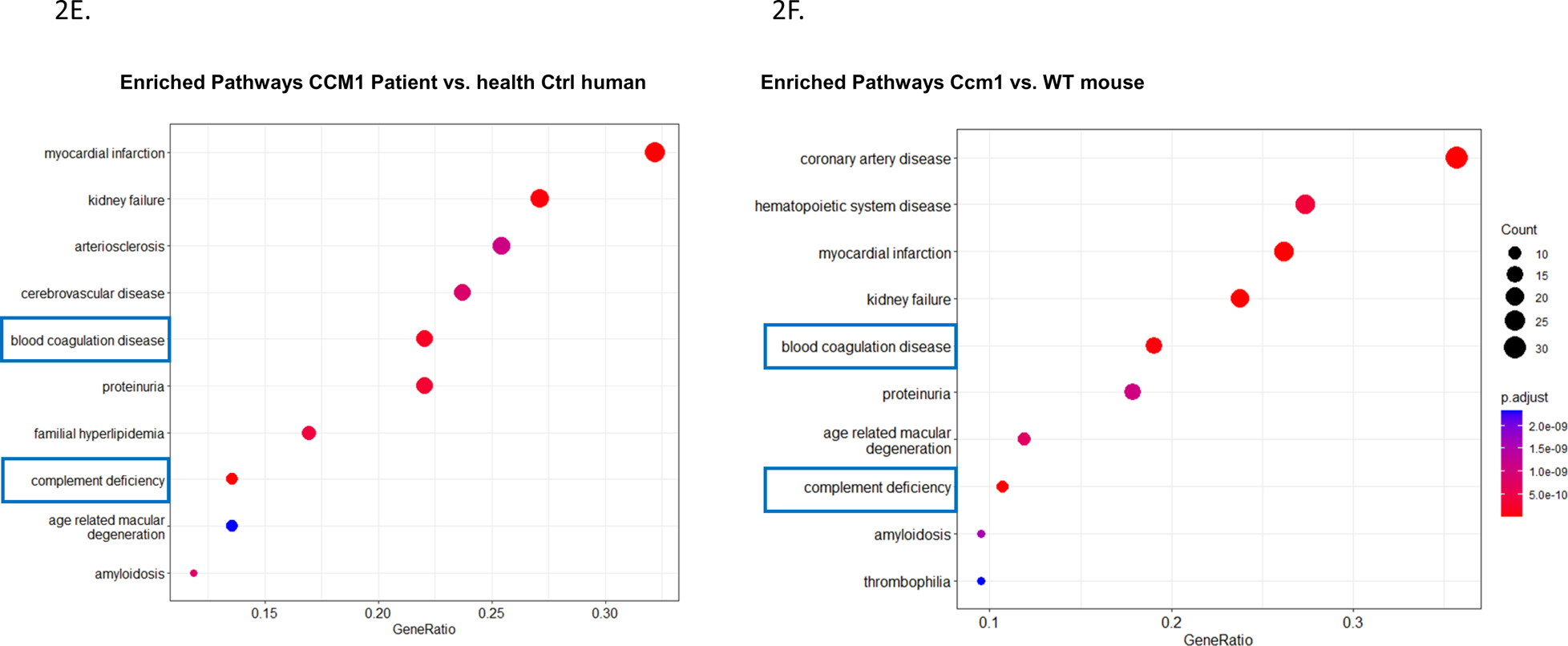
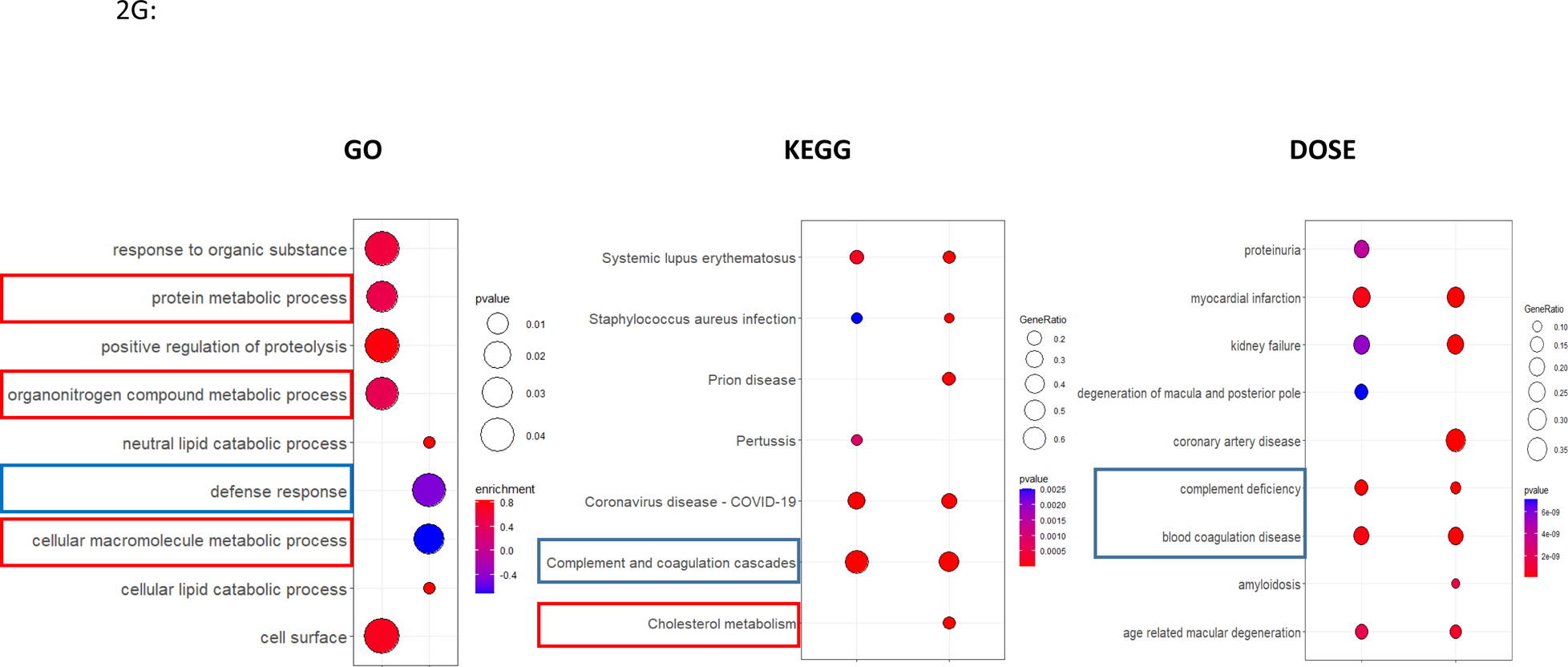
Comparative proteomics identify key pathways and their associated biomarkers for CCM1 deficiency. Comparative proteomics research has revealed crucial pathways and associated biomarkers for CCM1 deficiency. By detecting differentially expressed serum proteins (DEPs) in CCM1-deficient and normal subjects, the study successfully uncovered various pathways implicated in the CCM1 deficiency. This was achieved through gene set enrichment analysis (GSEA) of the identified DEPs in both human and mouse subjects. **A**-**B**. To assess the statistical significance of a biological process among the identified DEPs, Gene Ontology (GO) groupings for functional interaction networks and pathway analyses were conducted. This was done to evaluate the enrichment of differentially expressed serum proteins (DEPs) between CCM1-deficient and normal conditions in both humans (**A**) and mice (**B**). Two common pathways between 2A and 2B were identified: the cellular metabolic pathway and the coagulation and complement pathway. **C-D.** A comparative proteomics study was carried out between CCM1-deficient and normal conditions, employing disease ontology (DOSE) pathways to enrich differentially expressed serum proteins (DEPs) in humans (C) and mice (D). The pathways emphasized by colored frames represent the top-selected pathways. Two common pathways between 2A and 2B were identified: the cellular metabolic pathway and the coagulation and complement pathway. **E-F**. The Kyoto Encyclopedia of Genes and Genomes (KEGG) pathways were employed to enrich differentially expressed serum proteins (DEPs) between CCM1-deficient and normal conditions in humans (**E**) and mice (**F**). Two common pathways between 2A and 2B were identified: the cellular metabolic pathway and the coagulation and complement pathway. **G.** The application of three distinct enrichment libraries showcases pathways impacted by the CCM1-deficiency in comparison to the control for both species studied. From left to right, we observe enriched gene ontology (GO), Kyoto Encyclopedia of Genes and Genomes (KEGG), and Disease Ontology Semantic and Enrichment (DOSE) analyses, revealing affected pathways in metabolism (red outline) and complement and coagulation cascade pathways (blue outline). The highlighted pathways across two different species hold potential for the discovery of etiological and prognostic biomarkers for future validation analysis.

The GO enrichment pathway analysis showed that both human and mouse CCM1-deficient subjects shared metabolic processes and pathways compared to their respective control groups (Fig. 2A, 2B). In contrast, the GO-based GSEA identified shared coagulation and complement pathways in CCM1-deficient subjects relative to the control groups in both human (Table 1A, Suppl. Figs. 3A, 4A) and mice (Table 1B, Suppl. Figs. 3B, 4B). These differences could be attributed to the distinct bioinformatics approaches and databases used in each method.

**Table 1.**
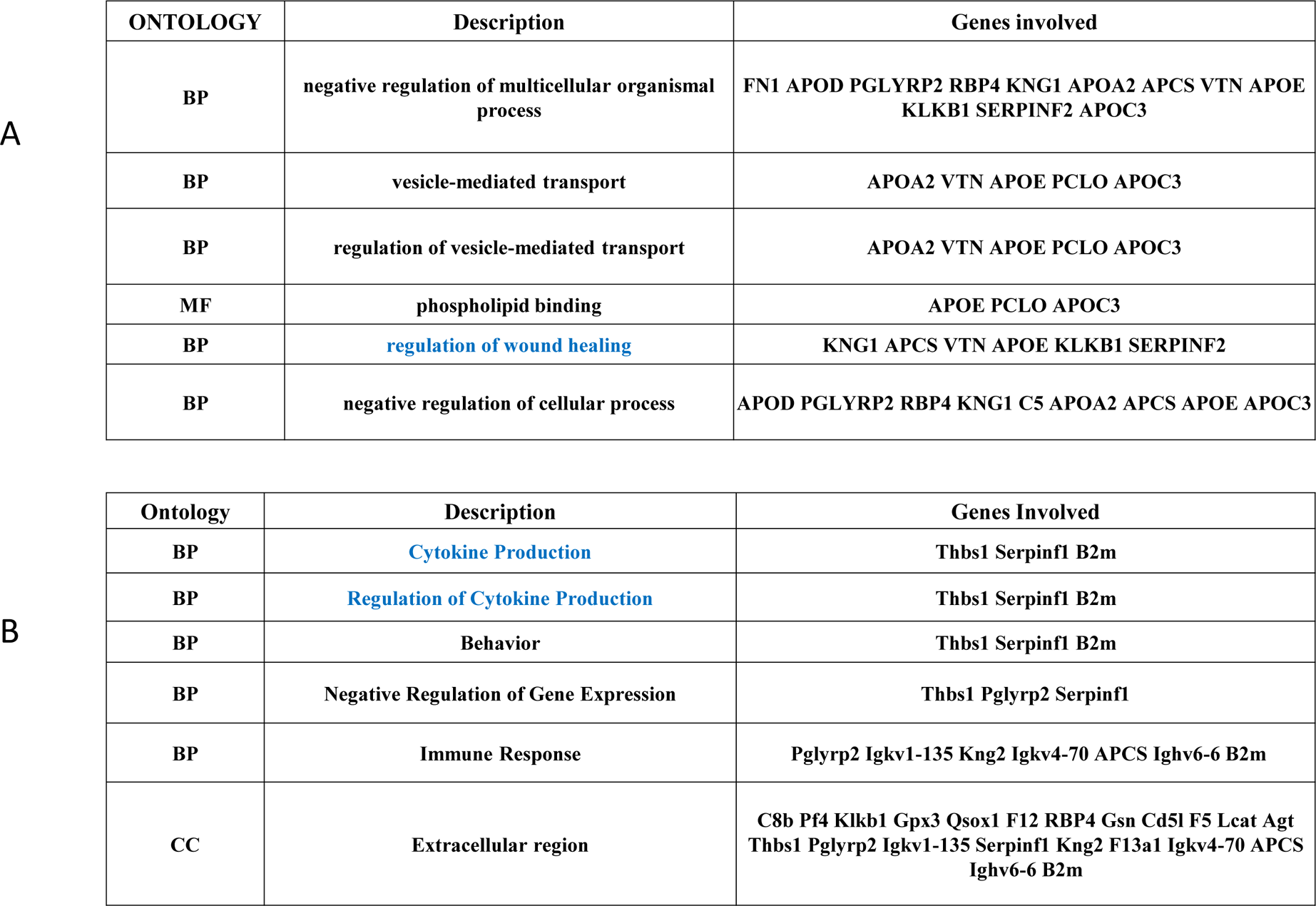
The gene set enrichment analysis (GSEA) on the GO. The gene set enrichment analysis (GSEA) was performed on the gene ontology (GO) of CCM1-deficient subjects compared to that of the control in both human (A) and mice (B). Significant DEP’s enrichment was then compared to a human database from BiocManager (Org.Hs.eg.db) (A) and to a mouse database from BiocManager (Org.Mm.eg.db) (B). The DEPs associated with pathways that showed significant differences in GO enrichment analysis are listed in the table. The pathways highlighted in blue represent the complement and coagulation cascade, with associated DEPs potentially serving as prognostic biomarkers.

### 3. Common shared CCM1-associated signaling pathways were identified in two species

To corroborate the GO enrichment findings, we subsequently performed disease ontology (DOSE) pathway analysis. This robust method facilitates the establishment of connections between enriched signaling pathways derived from differentially expressed genes and proteins (DEGs/DEPs) and the clinical phenotype (hemorrhagic CCMs), allowing for the identification of potential pathways linked to the progression of the pathology^50, 54, 56^.

It is noteworthy that the DOSE enrichment pathway analysis revealed the sharing of both metabolic processes and pathways and coagulation and complement pathways in individuals with CCM1 deficiency compared to the control group, in both humans and mice (as shown in Fig. 2C, 2D). However, DOSE-based GSEA analysis only identified coagulation and complement pathways (as demonstrated in Table 2A, 2B), possibly due to the reasons mentioned earlier. These results suggest that the differentially expressed proteins associated with metabolic processes/pathways and coagulation and complement pathways could have a crucial role in the development of CCM1 deficiency in humans and mice. In general, these findings provide further evidence supporting the involvement of proteins from these two significant pathways in CCM1 deficiency and emphasize their potential as biomarkers for this disease.

**Table 2.** The gene set enrichment analysis (GSEA) on the DOSE. The GSEA was performed on the disease ontology (DOSE) of CCM1-deficient subjects compared to that of the control in both human (A) and mice (B). Significant DEP’s enrichment was then compared to a human database from BiocManager (Org.Hs.eg.db) (A) and to a mouse database from BiocManager (Org.Mm.eg.db) (B). The DEPs associated with pathways that showed significant differences in GO enrichment analysis are listed in the table. The pathways highlighted in blue represent the complement and coagulation cascade, with associated DEPs potentially serving as prognostic biomarkers.

| DO:ID | Description | Gene Symbol |
| --- | --- | --- |
| DOID:6262 | Complement Deficiency | APOA1, C3 ,C4B, C5, C6, C7, C8A, C8B, CFH, CFHR1 |
| DOID:1247 | Blood Coagulation Disease | C3, C4B, C5, CFH, CPB2, F11, F13B, FGA, PROS1, SELL, SERPINA10, SERPIND1, SERPINF2, VWF |
| DOID:5844 | Myocardial Infarction | APOA1, APOA4, APOE, C3, C4B, C7, C8A, C8B, CFH, CPB2, F11, F13B, FGA, GSN, ITIH4, LPA, SERPIND1, VWF |
| DOID:10871 | Age Related Macular Degeneration | APOE, C3, C5, CFD, CFHR1, SELL, SERPINF1, TF |
| DO: 576 | Proteinuria | APOA1, APOA4, APOE, C3, C4B, C5,C6, CFH, CPB2, HPX, LPA, VTN |
| DO:1074 | Kidney Failure | AMBP, APOA1, APOA4, APOE, C3, C6, CFH, CPB2, FGA, PROS1, S100A8, SELL, SERPINF1, TFRC, VWF |

B
| DO: ID | Description | Gene Symbol |
| --- | --- | --- |
| DOID:3393 | Coronary Artery Disease | APOA1, APOA2, APOA4, APOC3, APOC4, APOE, APOM, C3, C7, C8A, C8B, CFH, CPB2, F11, F13B, F5, FGA, GSN, ITIH3, ITIH4, LCAT, LPA, LUM, PON3, SELL, SERPIND1, SERPINF2, SERPING1, VTN, VWF |
| DOID:626 | Complement Deficiency | APOA1, C3,C5, C6, C7, C8A, C8B, CFH, CFHR1 |
| DOID:1074 | Kidney Failure | AMBP, APOA1, APOA4, APOC3, APOE, B2M, C3, C6, CFH, CPB2, F5, FGA, LCAT, PROS1, PTGDS, S100A8, SELL, SERPINF1, TFRC, VWF |
| DOID:5844 | Myocardial Infarction | APOA1, APOA4, APOC3, APOE, C3, C7, C8A, C8B, CFH, CPB2, F11, F13B, F5, FGA, GSN, ITIH3, ITIH4, LCAT, LPA, SERPIND1, SERPING1, VWF |
| DOID:1247 | Blood Coagulation Disease | C3,C5, CFH, CP, CPB2, F11, F13B, F5, FGA, KLKB1, PROS1, SERPINA10, SERPIND1, SERPINF2, VWF |
| DOID:74 | Hematopoietic System Disease | APOE, C3,C5,CA1,CP, CPB2, F11, F13B, F5, FGA, KLKB1, MASP2, SELL, SERPINA10, SERPIND1, SERPINF1, SERPINF2, TF, TFRC, VWF |

In addition to GO and DOSE enrichment analyses, we also conducted an enriched KEGG analysis. This revealed that both human (Fig. 2E, Suppl. Fig. 5A) and mice (Fig. 2F, Suppl. Fig. 5B) CCM1-deficient subjects exhibited alterations in coagulation and complement pathways compared to the control group, which is further supported by KEGG-based GSEA (Table 3A, 3B). Notably, the top three pathways identified in the enriched KEGG analysis, including complement and coagulation, ECM-receptor interaction, and focal adhesion pathways, are all closely related to hemorrhagic events. These findings offer valuable insights into the biological processes underlying CCM1 deficiency and may contribute to the development of new therapeutic approaches for the disease.

**Table 3.** The gene set enrichment analysis (GSEA) on the KEGG. The GSEA was performed on the Kyoto Encyclopedia of Genes and Genomes (KEGG) pathways of CCM1-deficient subjects compared to that of the control in both human (A) and mice (B). Significant DEP’s enrichment was then compared to a human database from BiocManager (Org.Hs.eg.db) (A) and to a mouse database from BiocManager (Org.Mm.eg.db) (B). Blue-highlighted pathways represent the complement and coagulation cascade, with associated DEPs potentially serving as prognostic biomarkers. Furthermore, red-highlighted pathways pertain to the metabolic pathway, where related DEPs are intended for evaluation as potential etiological biomarkers.

| ID | Description | Gene Symbol |
| --- | --- | --- |
| hsa04610 | Complement and coagulation cascades | A2M, C1QA, C1QC, C3, C4BP, C6, C8A, F11, F13B, FGA, MASP2, SERPIND1, SERPINF2, VTN VWF |
| hsa05171 | Coronavirus disease - COVID-19 | C1QA, C1QC, C3, C6, C8A, F13B, FGA, MASP2, VWF |
| hsa05322 | Systemic lupus erythematosus | C1QA, C1QC, C3, C6, C8A, H2AJ |
| hsa05150 | Staphylococcus aureus infection | C1QA, C1QC, C3, KRT9, MASP2 |
| hsa05133 | Pertussis | C1QA, C1QC, C3, C4BPB |
| hsa04613 | Neutrophil extracellular trap formation | C3, FGA, H2AJ, VWF |
| hsa04512 | ECM-receptor interaction | CD44, VTN, VWF |
| hsa05142 | Chagas disease | C1QA, C1QC, C3 |

B
| ID | Description | Gene Symbol |
| --- | --- | --- |
| hsa04610 | Complement and coagulation cascades | C1QA, C1QB, C1QC, C3, C4BPB, C5, C6, C7, C8A, C8B, CFD, CFH, CFHR1, CPB2, F11, F13B, F5, FGA, KLKB1, MASP2, PROS1, SERPIND1, SERPINF2, SERPING1, VTN, VWF |
| hsa05171 | Coronavirus disease - COVID-19 | C1QA, C1QB, C1QX, C3, C5, C6, C7, C8A, C8B, CFD, F13B, FGA, IKBKG, MASP2,VWF |
| hsa05150 | Staphylococcus aureus infection | C1QA, C1QB, C1QC, C3, C5, CFD,CFH, KRT9, MASP2 |
| hsa04979 | Cholesterol metabolism | APOA1, APOA2, APOA4, APOC3, APOE, LCAT, LPA |
| hsa05322 | Systemic lupus erythematosus | C1QA, C1QB, C1QC, C3, C5, C6, C7, C8A, C8B, H2AJ |
| hsa05133 | Pertussis | C1QA, C1QB, C1QC, C3, C4BPB, C5, SERPING1 |

### 4. Candidate serum circulating biomarkers were identified through comparative analysis among common shared CCM1-associated signaling pathways in two species

Finally, we conducted a comprehensive comparative analysis among three separate enrichment approaches (GO/DOSE/KEGG) to identify pathways affected by CCM1-deficiency in comparison to the control group across both species. Our comparative pathway dotplots reveal that both metabolic processes/pathways and coagulation and complement pathways are consistently present in the serum of subjects with CCM1 deficiency relative to the control group, in both humans and mice (Fig. 2G). This observation is further substantiated by the GSEA results derived from the three distinct enrichment libraries (Table 4A, 4B). Our findings indicate that the identified serum DEPs mainly linked to two primary pathways could act as potential circulating blood biomarkers for the disease, namely metabolic processes/pathways and coagulation and complement pathways. Recent research in our lab has shown that CCM1 loss-of-function (LOF) results in disrupted metabolic processes/pathways, which aligns with prior studies highlighting the association between metabolic disturbances, oxidative stress, and intracellular reactive oxygen species (ROS) with CCM1 deficiency^57–60^. Additionally, we identified serum DEPs associated with coagulation and complement pathways, including complement C3, fibrinogen, vitronectin, collagen, fibronectin, and laminin. These findings align with recent studies,. In total, we discovered 21 potential blood biomarkers among the 71 detected DEPs. In sum, this study identified 21 potential blood biomarkers, primarily within metabolic processes/pathways and the complement and coagulation cascade pathway, which could serve as etiological or prognostic blood biomarkers, respectively. These biomarkers were found in both BACA CCM1 hemizygous mutation patients and Ccm1 hemizygous mutant mice using three distinct pathway enrichment methods with separate library datasets. The consistent results across approaches reinforce the findings and emphasize the potential of these biomarkers as diagnostic tools for CCM1 deficiency.

**Table 4.** The combined GSEA from three distinct enrichment libraries showcases pathways impacted by the CCM1-deficiency in comparison to the control for both species. Blue-highlighted pathways represent the complement and coagulation cascade, with associated DEPs potentially serving as prognostic biomarkers. Additionally, red-highlighted pathways pertain to the metabolic pathway, where related DEPs are intended for evaluation as potential etiological biomarkers.

| Gene symbol | Gene name | Pathway |
| --- | --- | --- |
| APCS | Amyloid P Component | complement and coagulation cascades |
| SERPINF1 | Serpin Family F Member 1 | complement and coagulation cascades |
| SERPINF2 | Serpin Family F Member 2 | complement and coagulation cascades |
| THBS1 | Thrombospondin 1 | complement and coagulation cascades |
| VTN | Vitronectin | complement and coagulation cascades |
| FN1 | Fibronectin 1 | complement and coagulation cascades |
| F12 | Coagulation Factor XII | complement and coagulation cascades |
| GSN | Gelsolin | complement and coagulation cascades |
| KLKB1 | Kallikrein B1 | complement and coagulation cascades |
| KNG1 | Kininogen 1 | complement and coagulation cascades |
| KNG2 | Kininogen 2 | complement and coagulation cascades |
| LCAT | Lecithin-Cholesterol Acyltransferase | cholesterol metabolism |
| APOA2 | Apolipoprotein A2 | lipoprotein metabolism |
| APOC3 | Apolipoprotein C3 | lipoprotein metabolism |
| APOD | Apolipoprotein D | lipoprotein metabolism |
| APOE | Apolipoprotein E | lipoprotein metabolism |
| PCLO | Piccolo Presynaptic Cytomatrix Protein | cytoskeletal matrix |
| QSOX1 | Quiescin Sulfhydryl Oxidase 1 | extracellular matrix |
| B2M | β2-Microglobulin | immuno pathways |
| RBP4 | Retinol Binding Protein 4 | membrane transporter |
| PGLYRP2 | Peptidoglycan Recognition Protein 2 | *N*-acetylmuramoyl-L-alanine amidase |

## Discussions

The objective of this research is to systematically detect prospective blood prognostic biomarkers in a homogeneous group of Hispanic fCCM patients and Ccm mutant mice, establishing a strong basis for our ongoing biomarker project. Our ongoing project aims to assess the potential of these biomarkers to predict early hemorrhagic events and recognize a critical time window for patients to receive preventive therapeutic treatment using deep learning algorithms combined with our candidate serum biomarkers. The proposed experiments will offer valuable insights into the development and prognosis of hemorrhagic CCMs and provide information about environmental exposures, effect modifiers, or risk factors linked to their progression. Gene Set Enrichment Analysis (GSEA) was used to identify signaling pathways enriched due to CCM1 deficiency. The top two pathways identified were metabolic processes/pathways and coagulation and complement pathways, which are relevant to hemorrhagic events. Coagulation signaling is linked to hemorrhagic stroke risk and outcomes^61–65^, leading to proposals for coagulation-targeted therapies and circulating prognostic biomarkers^62, 66^. Similarly, the complement cascade has also been associated with hemorrhagic stroke. However, it is still under debate whether complement factors can be used as prognostic tools for pre-hemorrhagic progression or as biomarkers for recovery after hemorrhagic events^61, 67–70^.

Among identified candidate prognostic biomarkers, plasma kallikrein (PKa) is involved in blood coagulation, fibrinolysis, hemostasis, and inflammatory response^71–74^. PKa deficiency due to KLKB1 mutations leads to vascular bleeding and has been implicated in hereditary angioedema and hemorrhagic stroke^73, 75–81^. Serpins, a superfamily of serine protease inhibitors, play key roles in vascular angiogenesis and have been implicated in retinal vascular leakage and hemorrhagic stroke ^82–89^. Peptidoglycan recognition protein 2 (PGLYRP2) is involved in immunomodulation and innate immunity, while the adenomatous polyposis coli (Apc) gene is crucial in development, negatively regulates Wnt signaling, and may be involved in angiogenesis ^90–95^. Retinol binding protein 4 (RBP4) is linked to the severity of cardiovascular disorders, and complement factors are known to be associated with hemorrhagic stroke ^61, 67–70, 96–99^. These biomarkers may help in understanding and treating various vascular conditions. There are several limitations to this study that should be acknowledged. Firstly, non-hemorrhagic CCMs (NHCs) were not included in the analysis due to the omics experimental design. Secondly, while this study has the largest sample size with both human and mice data, our power analysis indicates that with the current sample size, we may have missed some potential targeted proteins, leading to type-2 errors. To address this, we plan to examine 19 subjects and 38 controls with a 1:2 ratio to achieve 80% power to detect large differences with a 0.05 significance level. We aim to complete our analysis by the end of this year with an even larger sample size of 48 subjects and 96 controls, which will enable us to detect moderate differences between them.

## Conclusions

This research identified potential blood-based etiological and prognostic biomarkers for CCM1 deficiency in large, uniform Hispanic populations and Ccm1-deficient mice. Both human and mouse subjects exhibited differentially expressed serum proteins (DEPs) that participated in a range of biological functions. Three methods of enrichment analysis, which include Gene Ontology (GO), Disease Ontology (DOSE), and KEGG pathway analysis, were employed to investigate the functional traits and biological pathways linked to CCM1 deficiency-related DEPs. The two primary pathways, metabolic and coagulation/complement pathways, which include 21 candidate blood-derived biomarkers, were found to be strongly associated with hemorrhagic incidents.

## Ethics approval and consent to participate

### Ethics approval

This investigative research is part of the ongoing biomarker project known as “CCMs Among Hispanic Population Study Group (CHIPS).” The CHIPS project has been conducted according to the World Medical Association (WMA) Declaration of Helsinki and received approval from the TTUHSCEP-IRB committee on January 01, 2021 (E21010). This project is actively enrolling participants via ClinicalTrials.gov under the identifier NCT05148663 starting on December 8, 2021.

### Consent to participate

Participation in the study is voluntary, and those who do not fulfill the eligibility criteria are not allowed to join. The consent process with written consent form, which is mandatory for all eligible participants (or assent forms for underage patients and their guardians), takes place prior to the blood collection.

## Consent for publication

Not applicable

## Author contributions

Detailed in the CREDIT author contribution information on the online submission system.

## Acknowledgement

We wish to thank Muaz Bhalli, Alexander Le, Ofek Belkin, Mellisa Renteria, David Jang, Justin Aickareth, Victoria Reid, Majd Hawwar, Revathi Gnanasekaran, Nickolas Sanchez, Charlie Harvey, and Drexell Vincent at Texas Tech University Health Science Center El Paso (TTUHSCEP); and Moqsadur Rahman and Mahmud Shahriar Hossain at University of Texas El Paso (UTEP) for their technical help during the experiments.

## Availability of data and materials

Some essential analytical data were provided in supplemental materials in the online version of the journal, while all data submitted were comply with Institutional or Ethical Review Board requirements and applicable government regulations.

