## Supplementary material for "Blood prognostic biomarker signatures for hemorrhagic cerebral cavernous malformations (CCMs)": Suppl Materials

### ***Supplemental Materials***

***Shared co-first authorship**: Authors who have worked together and contributed equally to this project.

[++]All correspondence:
Jun Zhang, Sc.D., Ph.D.
Department of Molecular and Translational Medicine (MTM)
Texas Tech University Health Science Center El Paso
5001 El Paso Drive, El Paso, El Paso, TX 79905

### Supplemental Legends

**Supplemental Figure 1. A biological function analysis on the number of expressed proteins by CCM1 subjects compared to normal controls using group gene ontogeny (GO).**

**A-B.** Group Gene Ontology (GO) analysis and pathway analysis were performed for biological profiling and systematic functional annotation of the identified group of DEPs in order to determine the statistical significance of a biological process. These key processes indicate the presence of proteins, but their effect on the study subject in both human (**A**) and mouse (**B**) cannot be determined solely based on this information. Supplemental figure 2 provides similar trends in molecular function (Suppl. Fig. 2A, 2C) and cellular processes (Suppl. Fig. 2B, 2D).

**C-D.** To confirm whether our identified DEPs are frequently associated with specific biological functions, GO enrichment analysis was performed on three distinct subtypes of GO: biological process, molecular function, and cellular component. The molecular function and cellular component subtypes can be observed in supplemental figure 2. It's important to note that these key nodes or processes indicate the presence of proteins, but their positive or negative effect on both human (**C**) and mouse (**D**) study subjects cannot be determined solely based on this information.

**Supplemental Figure 2. Group Gene Ontology (GO) analysis on the molecular function and cellular component of the expressed proteins in both human and mouse models with CCM1 deficiency, compared to their respective healthy controls.**

**2A.** The group analysis of molecular function in the human CCM1 subject indicated that the majority of the expressed proteins were involved in binding processes. However, it's important to note that the number of proteins measured does not reflect whether these processes are positively or negatively regulated. Nonetheless, the higher number of proteins may suggest an increased demand due to decreased function observed in CCM1 diagnosis. **2B.** The cellular component analysis of the human CCM1 subject revealed that the majority of the expressed proteins are involved in cellular anatomical entity. It's important to note that the number of proteins produced does not indicate whether their effects are positive or negative. However, we can speculate that the increased demand for these proteins may be a result of the observed changes in CCM1 diagnosis.

**2C.** Similar to the human CCM1 subject (Supp. Fig. 2A), the molecular function group analysis of the mouse CCM1 subject showed that the largest number of expressed proteins were involved in binding actions. However, it's important to note that these numbers do not reflect whether the effects of these proteins are positive or negative. Nonetheless, the increased demand for these proteins could be due to the negative effects observed in CCM1 diagnosis. **2D.** Similarly, the cellular component group analysis of the mouse CCM1 subject showed that, like the affected human (Supp. Fig. 2B), most of the expressed proteins were involved in the cellular anatomical entity. **2E.** This panel depicts the pathway of protein activity within the molecular function pathways of a human CCM1 patient, with highlighted nodes indicating where the measured proteins from the collected samples interact with the pathway. It provides an expanded view of how the binding processes from Supp. Fig. 2A are affected. **2F.** This panel illustrates the pathway of protein activity within the cellular components of cells in a human CCM1 subject, highlighting the affected areas that support the CCM1 diagnosis, as well as where the abundant proteins from Supp. Fig. 2B are affecting the cellular components. **2G.** This molecular function pathway for the CCM1 mouse illustrates the areas where the harvested proteins affect the pathway, similar to what was observed in the human subject. The nodes represent the different stages of protein activity. As seen in both the mouse and human subjects, the binding activity is where these proteins are predominantly represented. Impaired binding processes are indicative of the CCM1 diagnosis, thus supporting the diagnosis in both laboratory and human subjects through replication. **2H.** This cellular component pathway for the CCM1 mouse subject illustrates the areas where the harvested proteins affect the cellular components. These areas, if negatively affected, would support the diagnosis of CCM1, as they are the impaired areas anticipated by the cellular malformations observed in CCM1 diagnosis. The nodes represent different stages of protein activity within the pathway.

**Supplemental Figure 3. GSEA analysis demonstrates the pathways affected in both human and mouse CCM1 subjects.**

**3A**. The gene set enrichment analysis upset plots illustrate the pathways that are affected in the CCM1 human subject. The affected pathways support the diagnosis of CCM1 subjects by showing negative regulation of organismal processes, negative responses to binding of cells as observed in wound healing, and negative effects on the organization of cellular processes. This is consistent with the biological process group GO as shown in supplemental figures 1A. **3B**. This gene set enrichment analysis upset plot displays the pathways that are affected in the mouse CCM1 subject. Similar to the human subject, there is a decreased function of the membrane, which supports the diagnosis of CCM1 as the malformation of the plasma membrane results in oblong formations observed in this diagnosis.

**Supplemental Figure 4: GSEA analysis identified enriched pathways for the DEPs in the CCM1-deficiencys compared to their normal conditions, for both human and mouse subjects.**

**4A**. The pathways identified in the human CCM1 subjects were compared to age and gender-matched healthy controls, revealing negative regulation of all enriched pathways that are known to be affected by CCM1 deficiency. This supports the previous upset plots that displayed the negatively affected pathways, as well as providing additional insight into the results from the group GO molecular function and cellular components observed in Supp. Fig. 1A-B. **4B**. The pathways identified through GSEA analysis of CCM1 mouse compared to a healthy control showed negative effects that signify the affected processes and further support the CCM1 diagnosis.

**Supplement Figure 5. Enriched KEGG pathways reaffirm the existence of complement and coagulation cascades caused by CCM1 deficiency in both Human and Mouse subjects.**

**5A**. The KEGG enrichment pathways for the Human enriched genome shows the complement and coagulation, including ECM components. While this was identified as a KEGG pathway, it in the human CCM1 patient it supports the degraded binding suggested by the GO enrichment where binding proteins were found (Supp. Fig. 2A). **5B. S**imilarly, the KEGG enrichment pathway analysis for the mouse Ccm1 subject highlighted the affected complement and coagulation cascades. This further supports our previous findings and strengthens the diagnosis of CCM1.

1-A.

1-B.

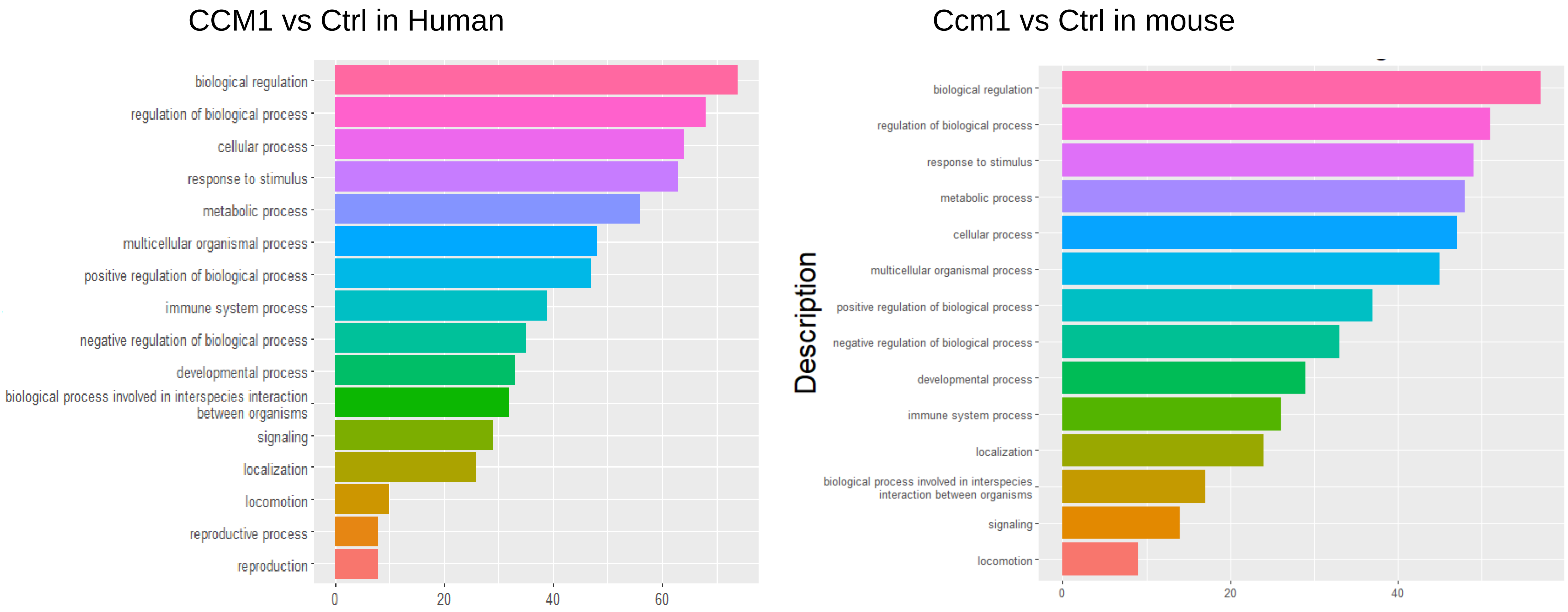


**1. Group-GO Biological Process**

1-C. Human

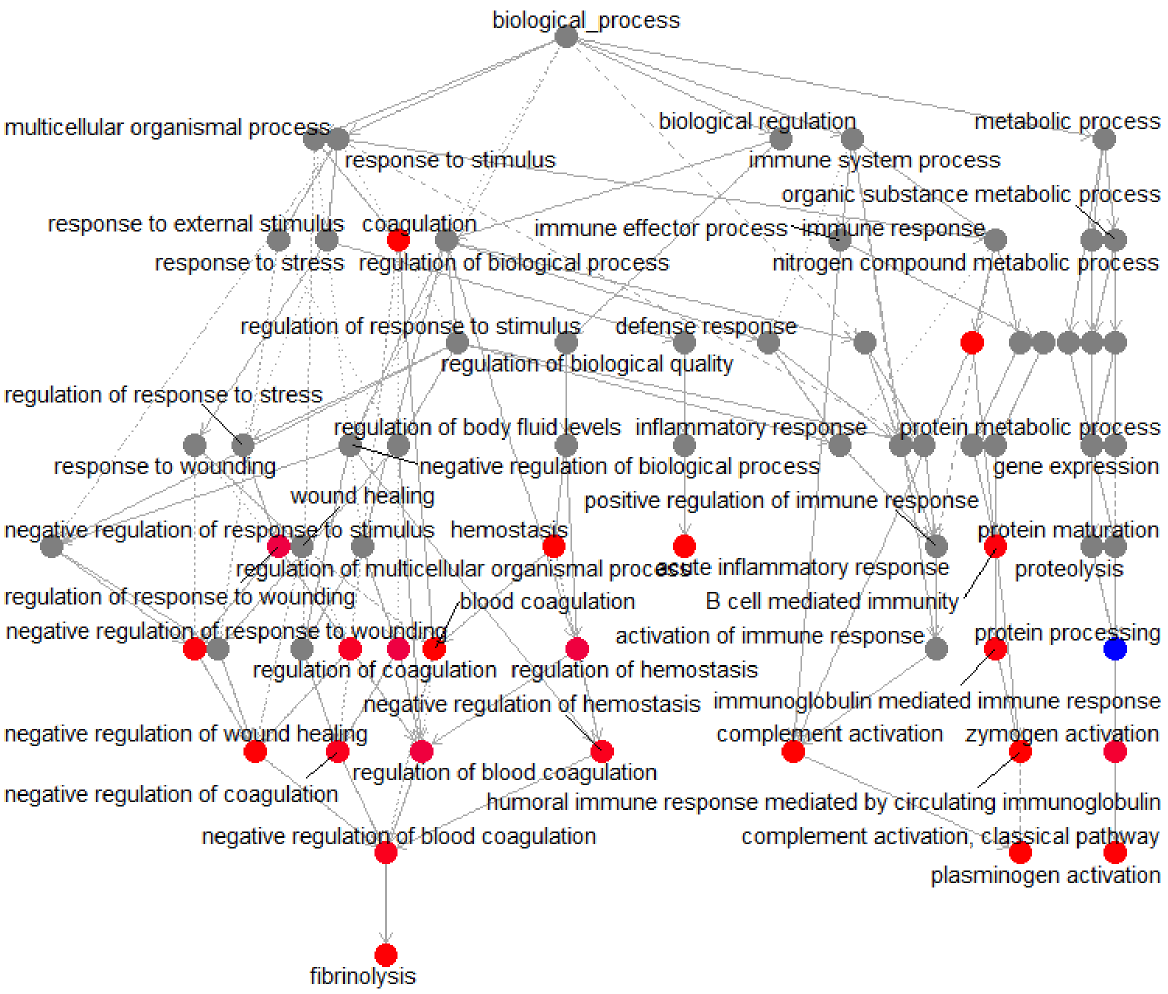


1-D. Mouse

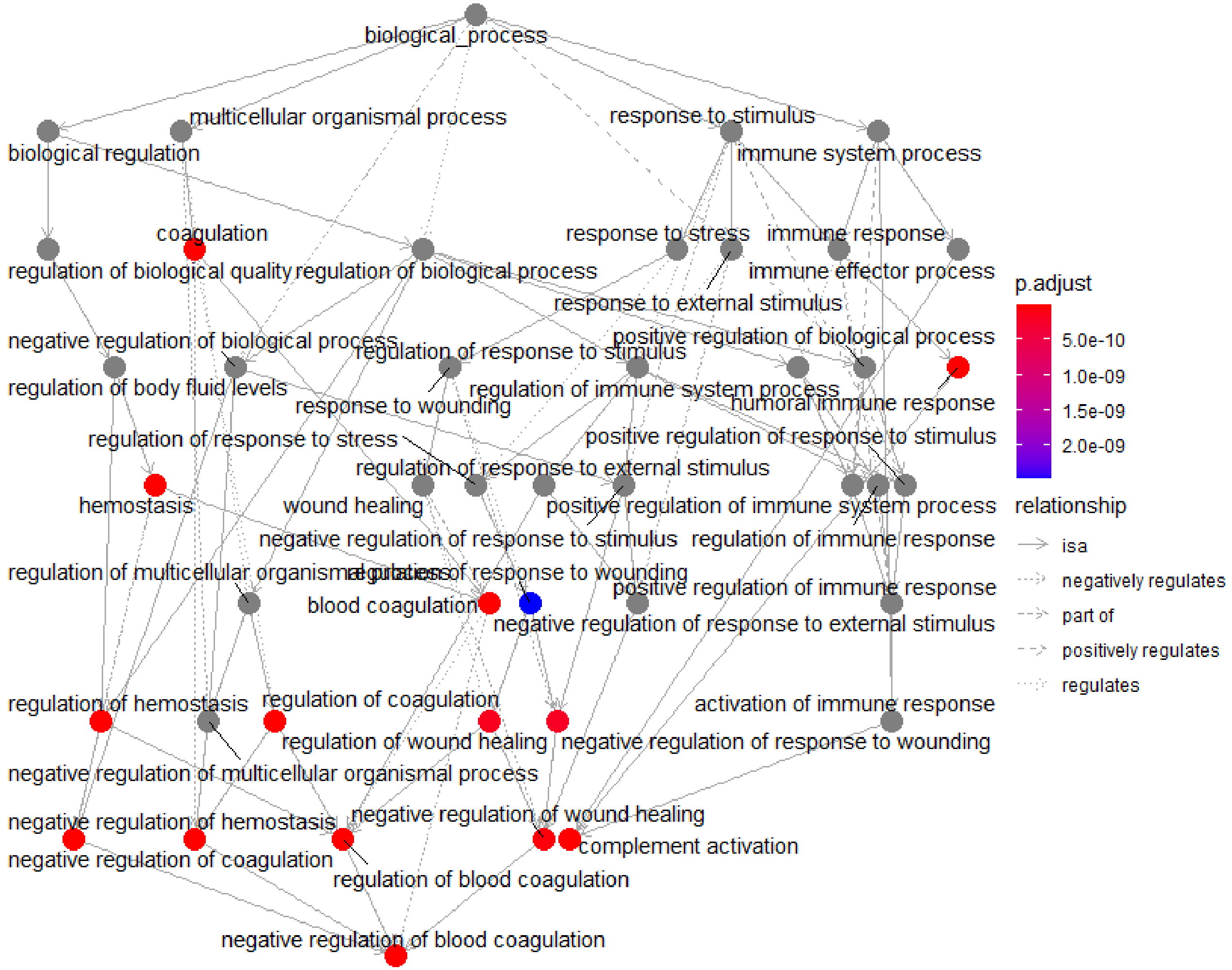


Enriched GO-Biological Process

2A.

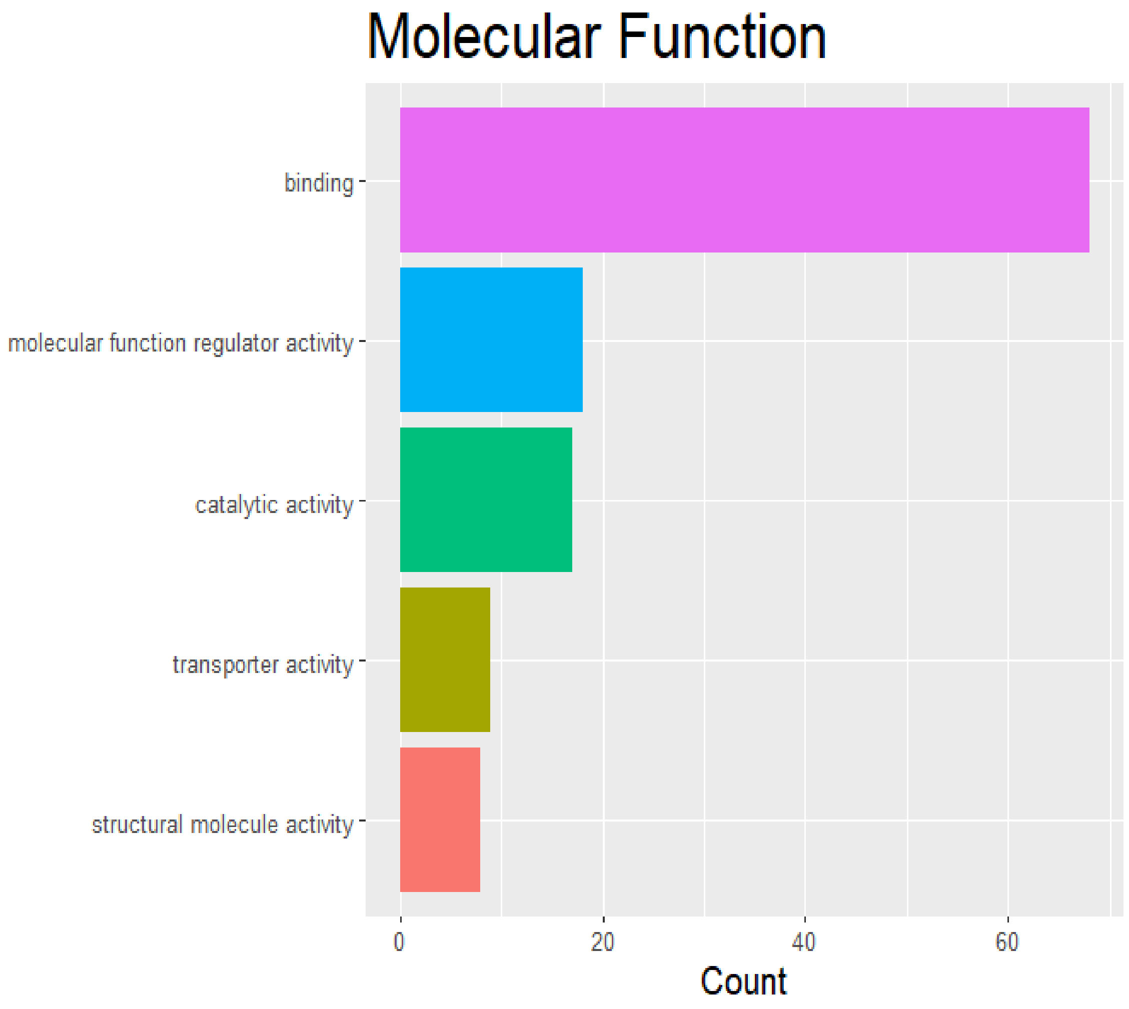

Molecular Function
binding
molecular function regulator activity
catalytic activity
transporter activity
structural molecule activity
0
20
40
60
Count


2B.

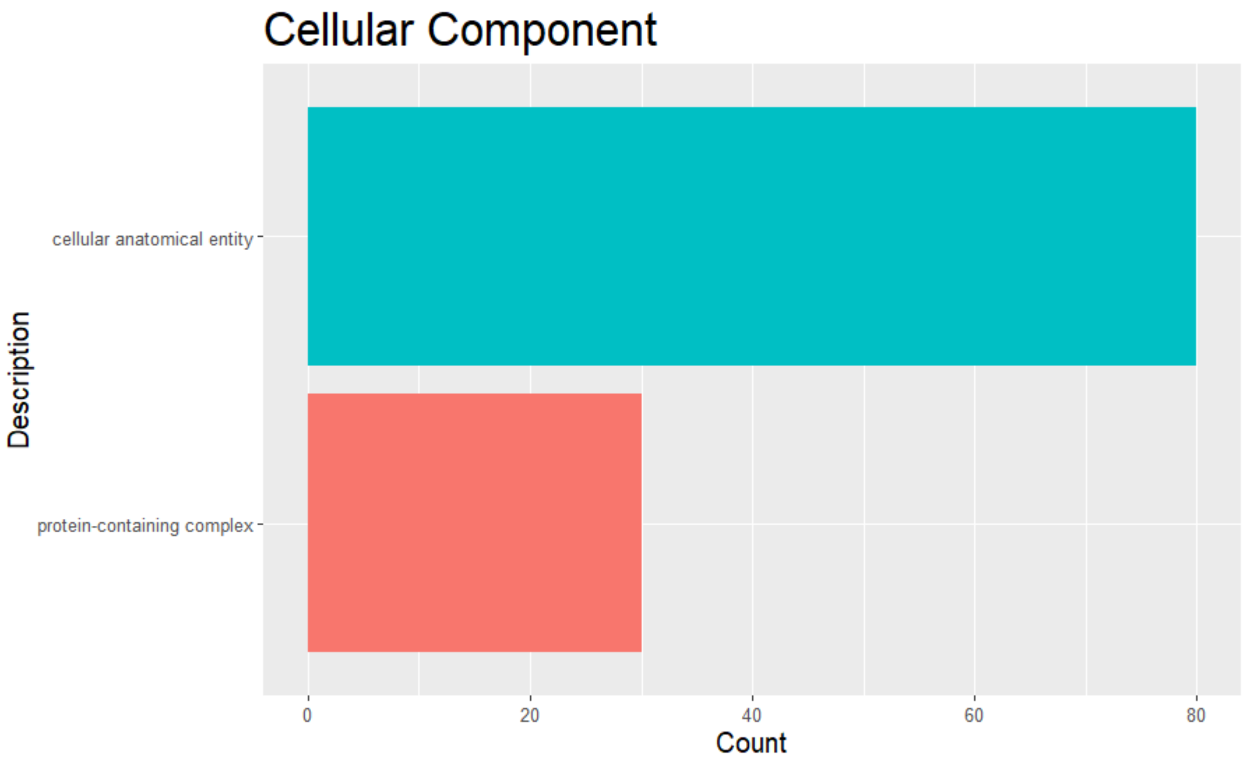

Cellular Component
Description
cellular anatomical entity
protein-containing complex
0
20
40
60
80
Count

2C.

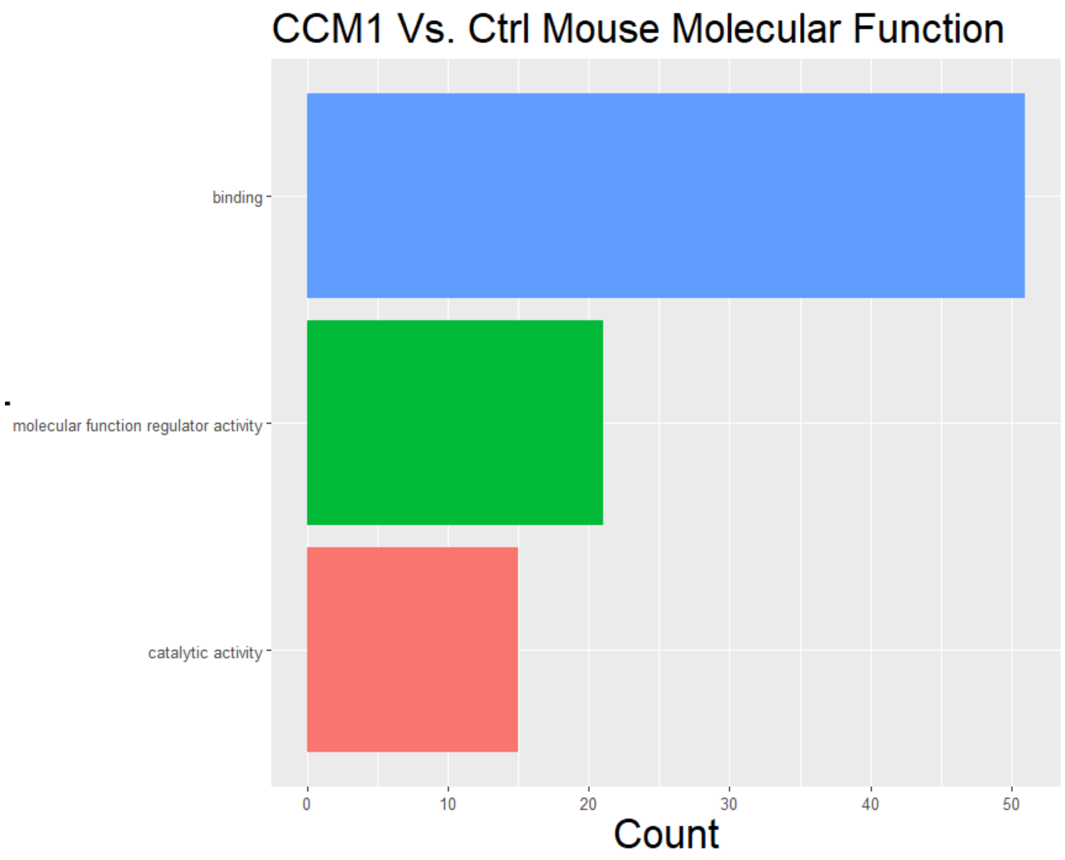
CCM1 Vs. Ctrl Mouse Molecular Function
binding
molecular function regulator activity
catalytic activity
0
10
20
30
40
50
Count

2D.

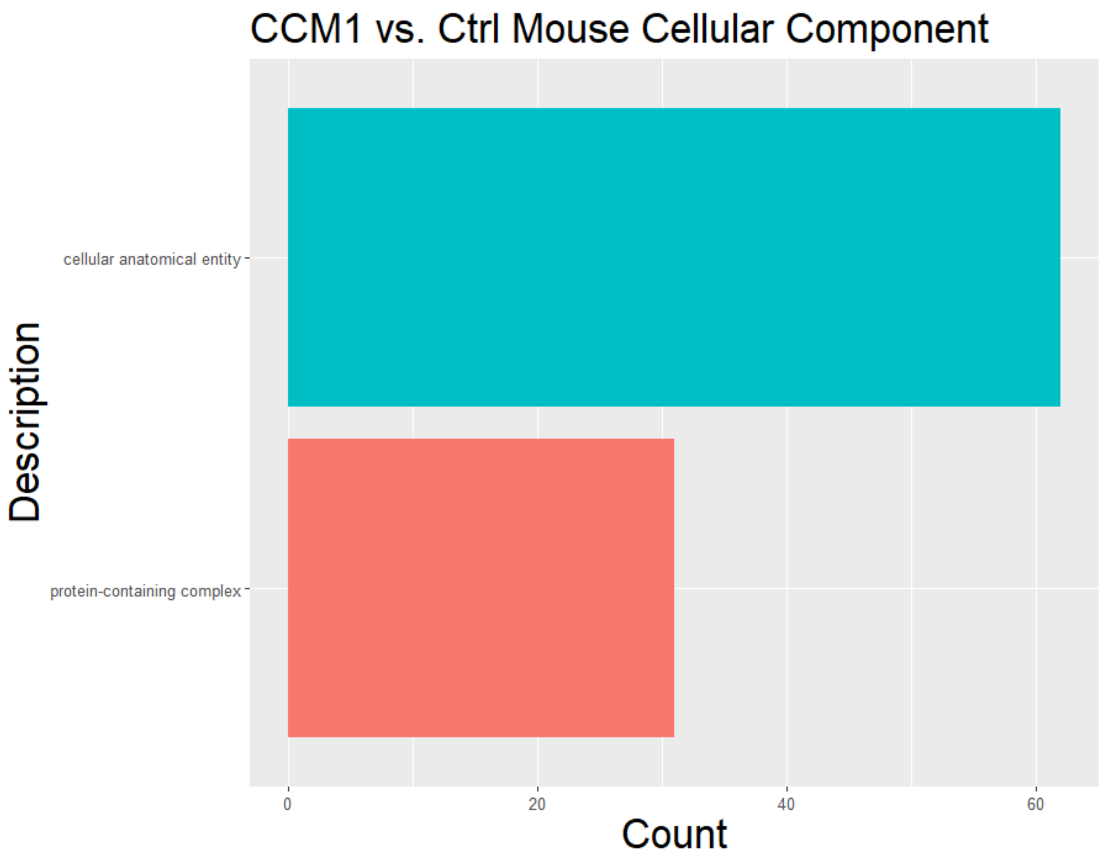
CCM1 vs. Ctrl Mouse Cellular Component
Description
cellular anatomical entity
protein-containing complex
0
20
40
60
Count

2E. human

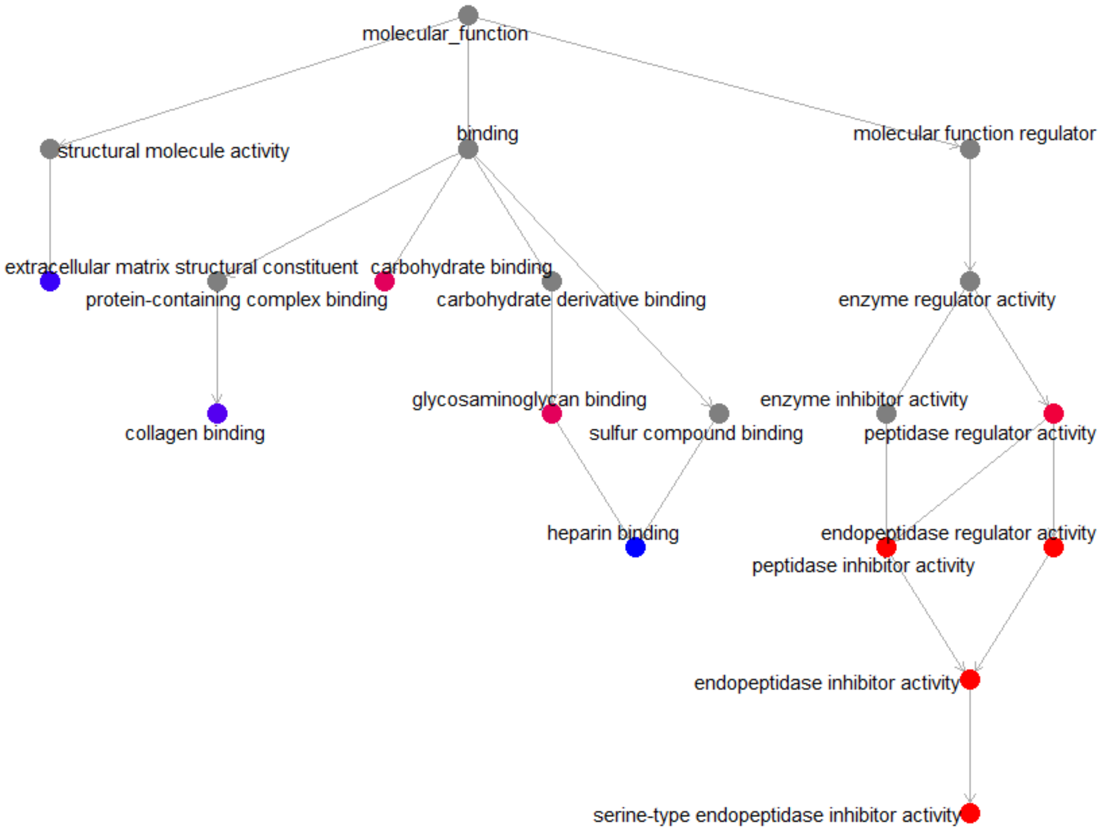
molecular_function
structural molecule activity
binding
molecular function regulator
extracellular matrix structural constituent
carbohydrate binding
protein-containing complex binding
carbohydrate derivative binding
enzyme regulator activity
collagen binding
glycosaminoglycan binding
enzyme inhibitor activity
sulfur compound binding
peptidase regulator activity
heparin binding
endopeptidase regulator activity
peptidase inhibitor activity
endopeptidase inhibitor activity
serine-type endopeptidase inhibitor activity

2F.

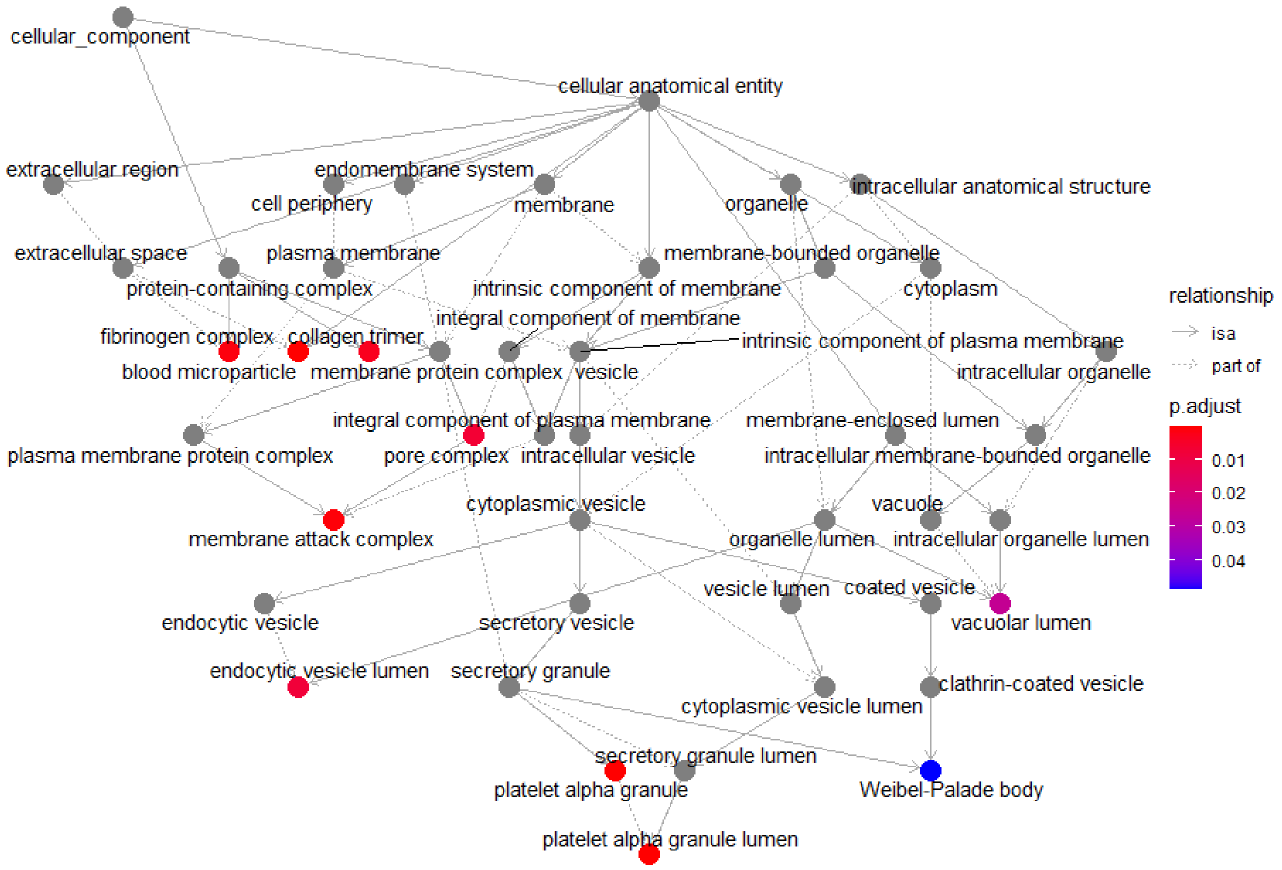
cellular_component
cellular anatomical entity
extracellular region
endomembrane system
cell periphery
membrane
organelle
intracellular anatomical structure
extracellular space
plasma membrane
membrane-bounded organelle
protein-containing complex
intrinsic component of membrane
cytoplasm
integral component of membrane
fibrinogen complex
collagen trimer
intrinsic component of plasma membrane
blood microparticle
membrane protein complex
vesicle
intracellular organelle
integral component of plasma membrane
membrane-enclosed lumen
plasma membrane protein complex
pore complex
intracellular vesicle
intracellular membrane-bounded organelle
cytoplasmic vesicle
vacuole
membrane attack complex
organelle lumen
intracellular organelle lumen
vesicle lumen
coated vesicle
endocytic vesicle
secretory vesicle
vacuolar lumen
endocytic vesicle lumen
secretory granule
clathrin-coated vesicle
cytoplasmic vesicle lumen
secretory granule lumen
platelet alpha granule
Weibel-Palade body
platelet alpha granule lumen
relationship
is a
part of
p.adjust
0.01
0.02
0.03
0.04

2G. Mouse

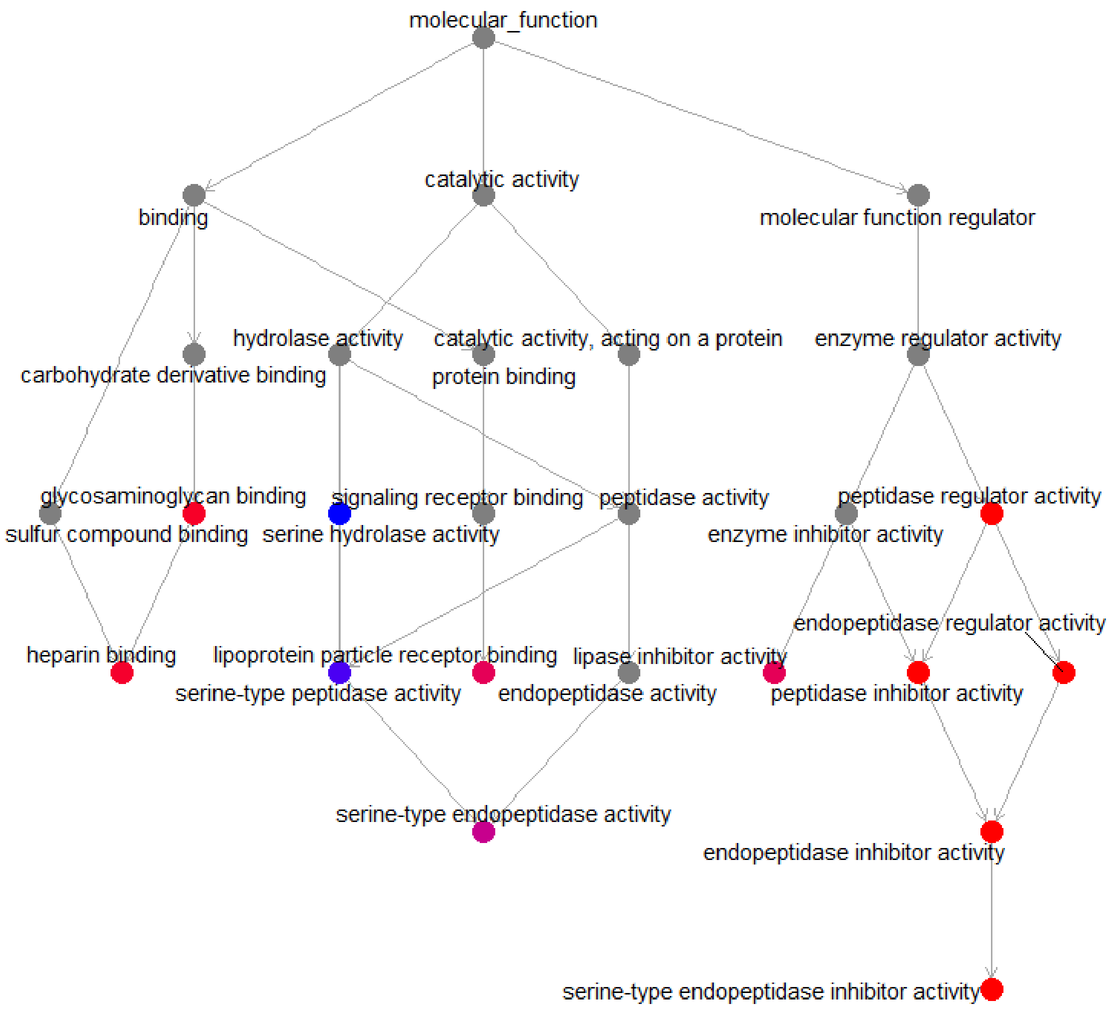
molecular_function
binding
catalytic activity
molecular function regulator
hydrolase activity
catalytic activity, acting on a protein
enzyme regulator activity
carbohydrate derivative binding
protein binding
glycosaminoglycan binding
signaling receptor binding
peptidase activity
peptidase regulator activity
sulfur compound binding
serine hydrolase activity
enzyme inhibitor activity
endopeptidase regulator activity
heparin binding
lipoprotein particle receptor binding
lipase inhibitor activity
serine-type peptidase activity
endopeptidase activity
peptidase inhibitor activity
serine-type endopeptidase activity
endopeptidase inhibitor activity
serine-type endopeptidase inhibitor activity

2H.

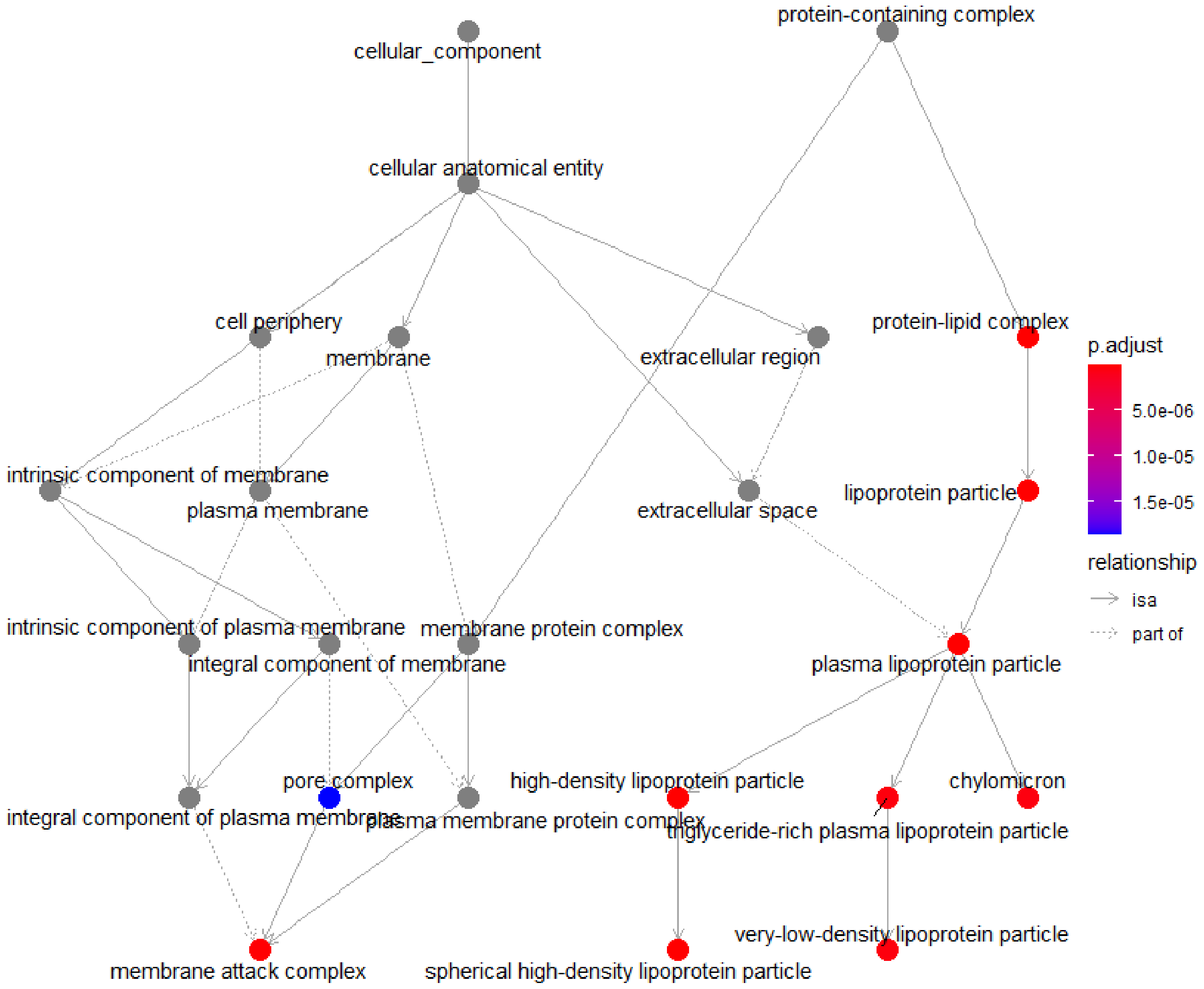
protein-containing complex
cellular_component
cellular anatomical entity
cell periphery
membrane
extracellular region
protein-lipid complex
p.adjust
5.0e-06
1.0e-05
1.5e-05
relationship
isa
part of
intrinsic component of membrane
plasma membrane
extracellular space
lipoprotein particle
intrinsic component of plasma membrane
membrane protein complex
integral component of membrane
plasma lipoprotein particle
pore complex
high-density lipoprotein particle
chylomicron
integral component of plasma membrane
plasma membrane protein complex
triglyceride-rich plasma lipoprotein particle
very-low-density lipoprotein particle
membrane attack complex
spherical high-density lipoprotein particle

3A.

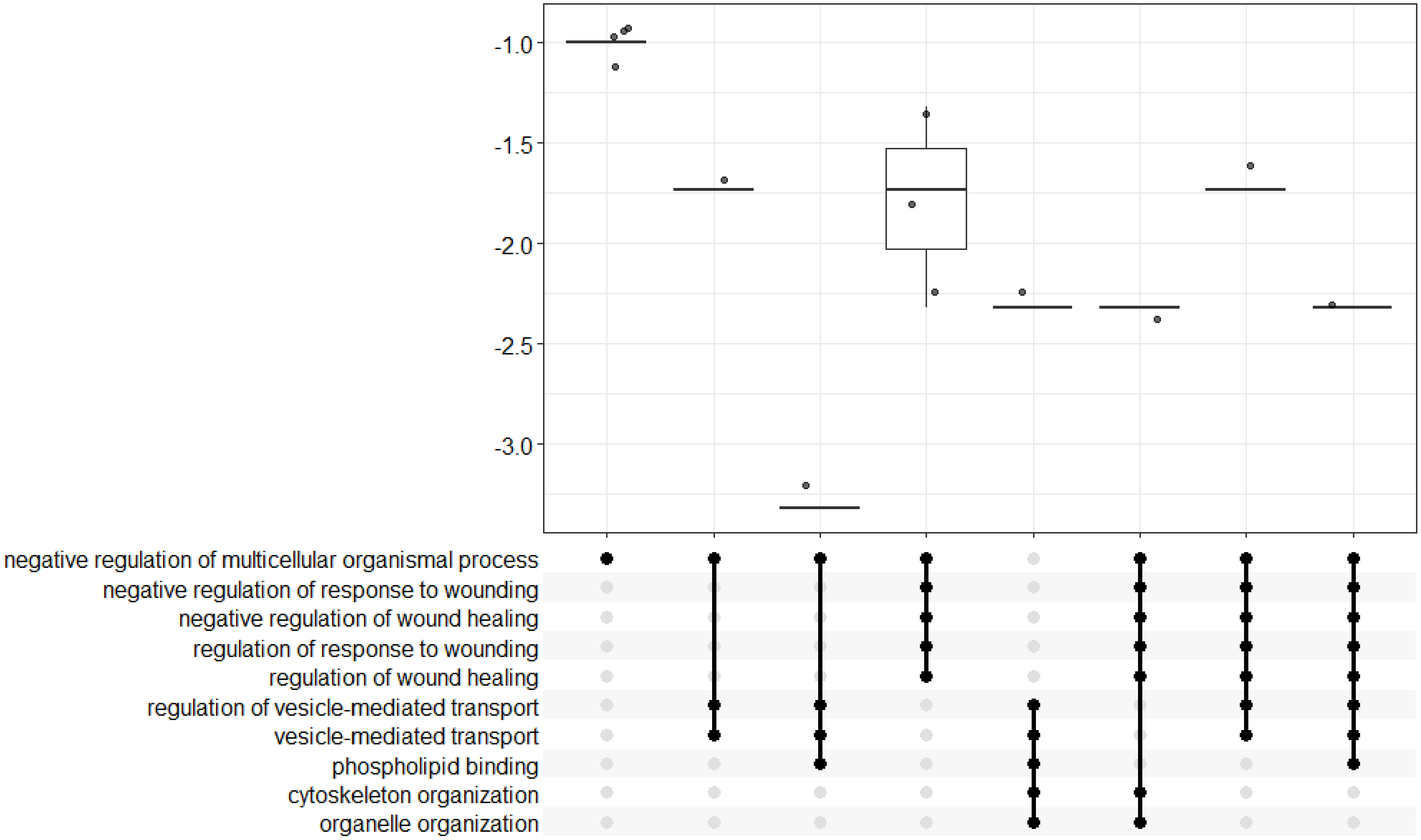

-1.0
-1.5
-2.0
-2.5
-3.0
negative regulation of multicellular organismal process
negative regulation of response to wounding
negative regulation of wound healing
regulation of response to wounding
regulation of wound healing
regulation of vesicle-mediated transport
vesicle-mediated transport
phospholipid binding
cytoskeleton organization
organelle organization

3B

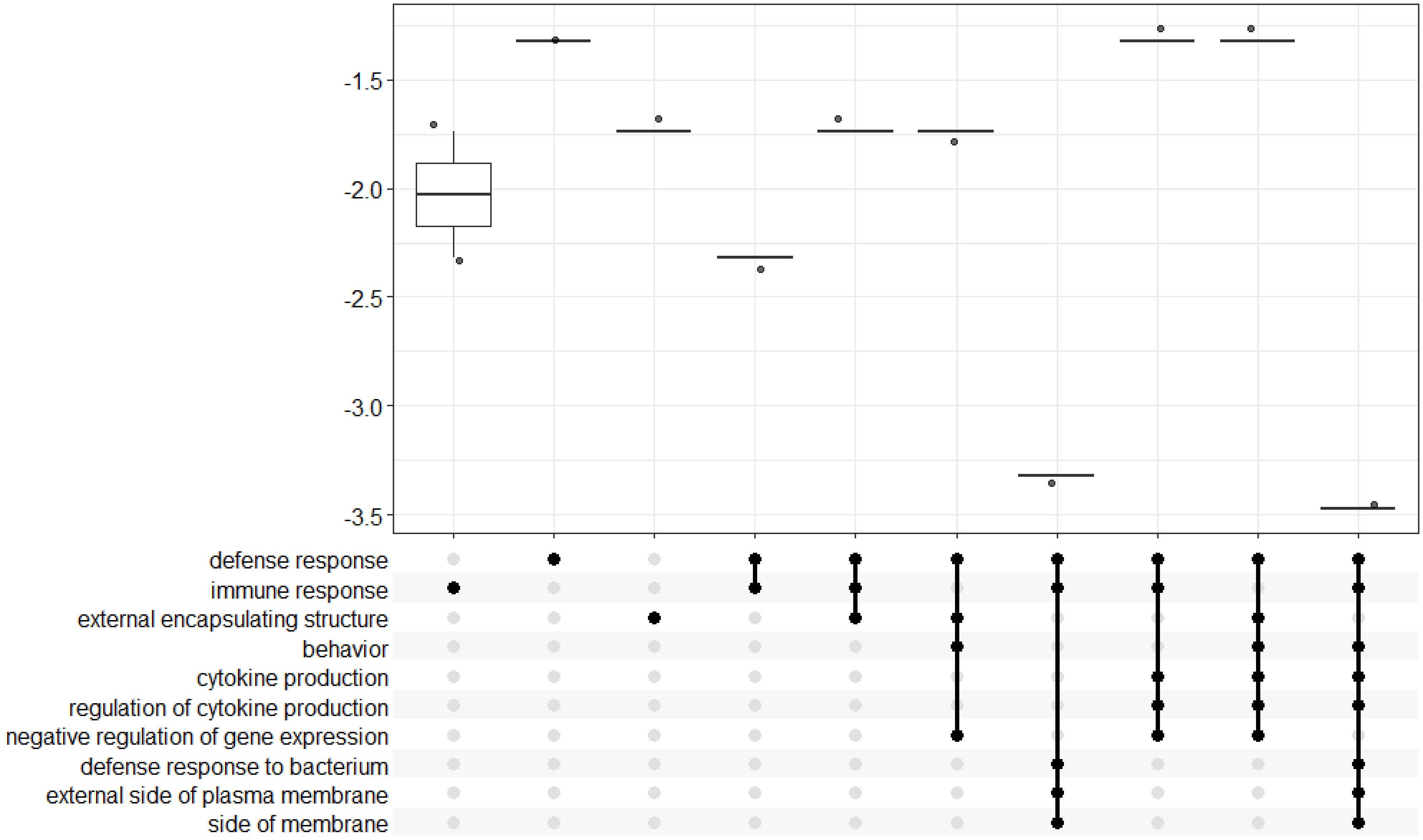
-1.5
-2.0
-2.5
-3.0
-3.5
defense response
immune response
external encapsulating structure
behavior
cytokine production
regulation of cytokine production
negative regulation of gene expression
defense response to bacterium
external side of plasma membrane
side of membrane

4A.

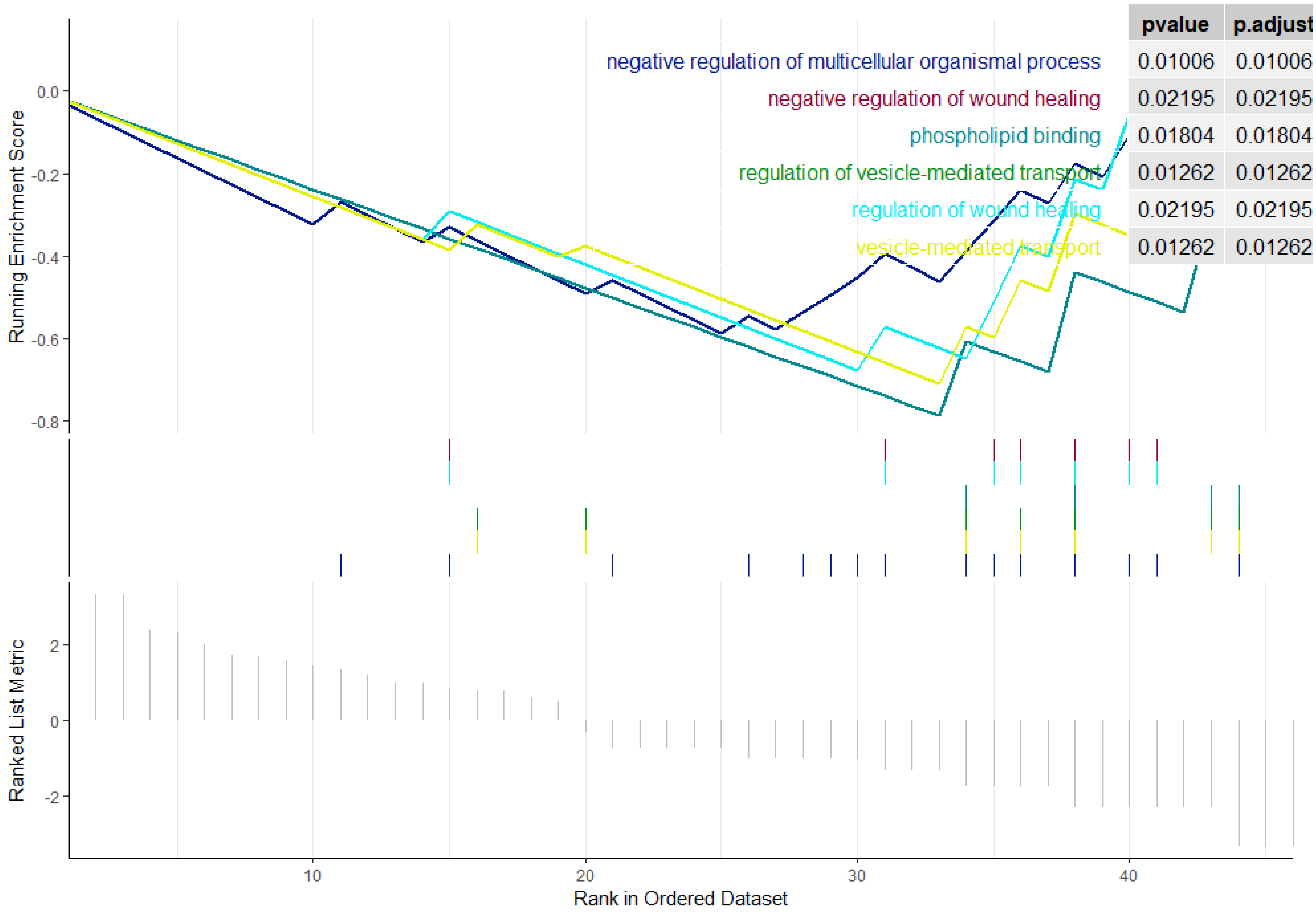
pvalue
p.adjust
negative regulation of multicellular organismal process
0.01006
0.01006
negative regulation of wound healing
0.02195
0.02195
phospholipid binding
0.01804
0.01804
regulation of vesicle-mediated transport
0.01262
0.01262
regulation of wound healing
0.02195
0.02195
vesicle-mediated transport
0.01262
0.01262
Running Enrichment Score
0.0
-0.2
-0.4
-0.6
-0.8
Ranked List Metric
2
0
-2
10
20
30
40
Rank in Ordered Dataset

4B.

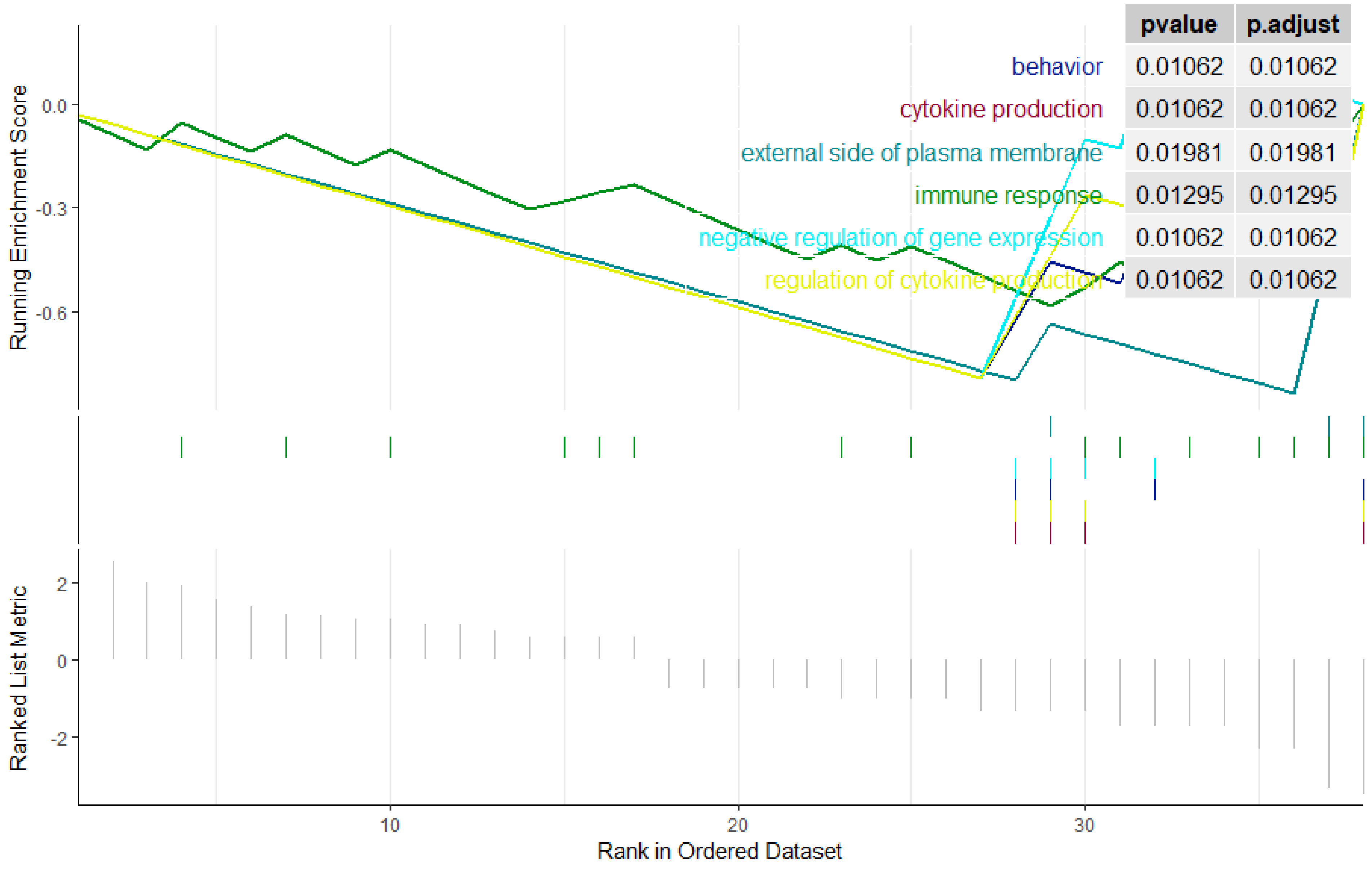
pvalue
p.adjust
behavior 0.01062 0.01062
cytokine production 0.01062 0.01062
external side of plasma membrane 0.01981 0.01981
immune response 0.01295 0.01295
negative regulation of gene expression 0.01062 0.01062
regulation of cytokine production 0.01062 0.01062
Running Enrichment Score
0.0
-0.3
-0.6
Ranked List Metric
2
0
-2
10
20
30
Rank in Ordered Dataset

5A.

Human KEGG Enriched pathway

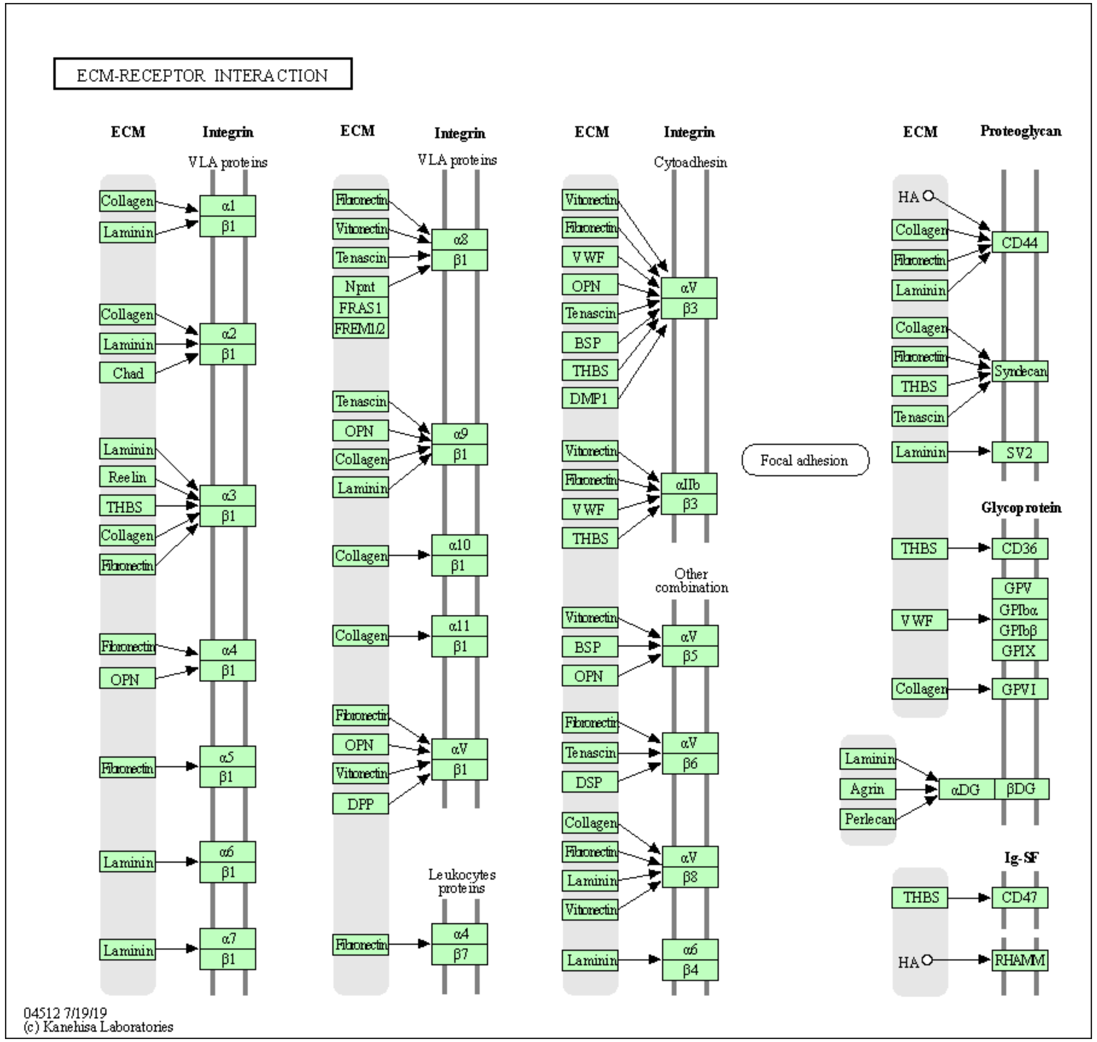


5B.

Mouse KEGG Enriched Pathway

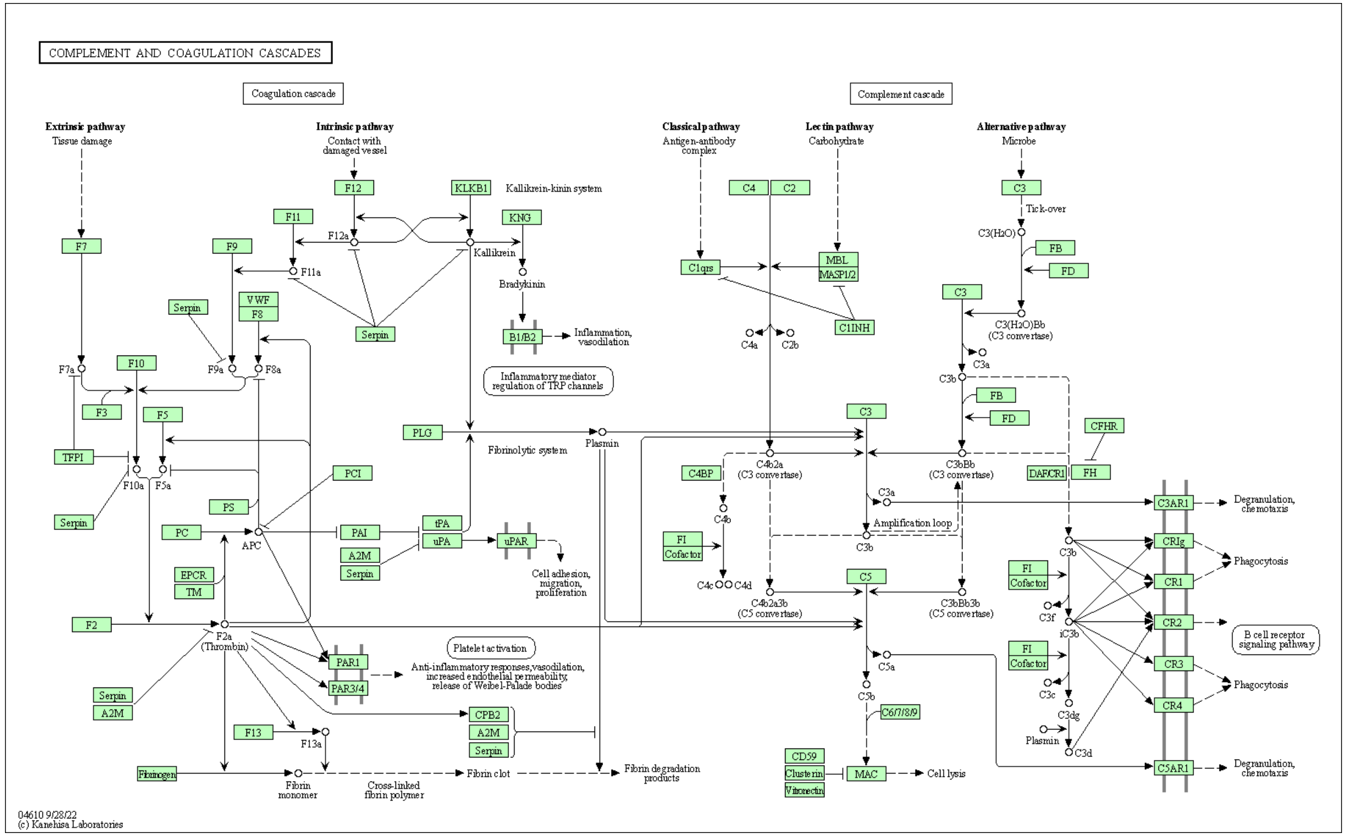
